# Defined human tri-lineage brain microtissues

**DOI:** 10.1101/2025.08.05.668605

**Authors:** Takeshi Uenaka, Sascha Jung, Ishan Kumar, Kit Vodehnal, Mohit Rastogi, Yongjin Yoo, Mark Koontz, Christian Thome, Wanhua Li, Tamara Chan, Erin M. Green, Kirill Chesnov, Zijun Sun, Shuyuan Zhang, Jinzhao Wang, Anthony Venida, Anne-Laure Mahul Mellier, Micaiah Atkins, Meredith Jackrel, Jan M. Skotheim, Tony Wyss-Coray, Monther Abu-Remaileh, Hilal A. Lashuel, Michael C. Bassik, Thomas C. Südhof, Antonio del Sol, Erik Ullian, Marius Wernig

**Affiliations:** Institute for Stem Cell Biology & Regenerative Medicine, Stanford University School of Medicine, Stanford, CA, 94305, USA; Department of Pathology, Stanford University School of Medicine, Stanford, CA 94305, USA; CIC bioGUNE-BRTA (Basque Research and Technology Alliance), Bizkaia Technology Park, 801 Building, Derio, 48160 Spain; Department of Ophthalmology, School of Medicine, University of California, San Francisco, San Francisco, CA, USA; Department of Molecular and Cellular Physiology, Stanford University School of Medicine, Stanford, CA 94305, USA; Department of Biological Sciences, University of Maryland Baltimore County, Baltimore, MD 21250, USA; Howard Hughes Medical Institute, Stanford University School of Medicine, Stanford, CA, USA; Department of Biology, Stanford University, Stanford, CA, USA; Department of Genetics, Stanford University, Stanford, CA, USA; Sarafan ChEM-H, Stanford University, Stanford, CA, USA; Laboratory of Molecular and Chemical Biology of Neurodegeneration, École Polytechnique Fédérale de Lausanne (EPFL), Lausanne, Switzerland; Department of Neurology and Neurological Sciences, Stanford University School of Medicine, CA, USA; Wu Tsai Neurosciences Institute, Stanford University School of Medicine, CA, USA; Department of Chemistry, Washington University in St. Louis, St. Louis, MO, USA; Chan Zuckerberg Biohub San Francisco, Stanford University, Stanford, CA, USA; Department of Chemical Engineering, Stanford University, Stanford, CA, USA; The Phil & Penny Knight Initiative for Brain Resilience at the Wu Tsai Neurosciences Institute, Stanford University, Stanford, CA, USA; Luxembourg Centre for Systems Biomedicine (LCSB), University of Luxembourg, 6 Avenue du Swing, Esch-Belval Esch-sur-Alzette, 4367 Luxembourg; IKERBASQUE, Basque Foundation for Science, Bilbao, 48013 Spain; Department of Chemical and Systems Biology, Stanford University School of Medicine, Stanford, CA 94305, USA

## Abstract

Microglia are the immune cells of the central nervous system and are thought to be key players in both physiological and disease conditions. Several microglial features are poorly conserved between mice and human, such as the function of the neurodegeneration-associated immune receptor Trem2. Induced pluripotent stem cell (iPSC)-derived microglia offer a powerful opportunity to generate and study human microglia. However, human iPSC-derived microglia often exhibit activated phenotypes *in vitro*, and assessing their impact on other brain cell types remains challenging due to limitations in current co-culture systems. Here, we developed fully defined brain microtissues, composed of human iPSC-derived neurons, astrocytes, and microglia, co-cultured in 2D or 3D formats. Our microtissues are stable and self-sufficient over time, requiring no exogenous cytokines or growth factors. All three cell types exhibit morphologies characteristic of their *in vivo* environment and show functional properties. Co-cultured microglia develop more homeostatic phenotypes compared to microglia exposed to exogenous cytokines. Hence, these tri-cultures provide a unique approach to investigate cell-cell interactions between brain cell types. We found that astrocytes and not neurons are sufficient for microglial survival and maturation, and that astrocyte-derived M-CSF is essential for microglial survival. Single-cell and single-nucleus RNA sequencing analyses nominated a network of reciprocal communication between cell types. Brain microtissues faithfully recapitulated pathogenic α-synuclein seeding and aggregation, suggesting their usefulness as human cell models to study not only normal but also pathological cell biological processes.

## INTRODUCTION

Human induced pluripotent stem cells (iPSCs) and embryonic stem cells (ESCs) have become valuable tools to investigate disease mechanisms because they can be differentiated into various somatic cell types, including those of the central nervous system (CNS), which are difficult to obtain from primary sources. Much attention has been dedicated to developing differentiation protocols to specific neuronal subtypes^1,2^ as those are the cells that are affected in many neurological diseases and they are not regenerated once lost. Although neurons can be cultured and survive for extended periods of time in defined media, physiologically, they do not exist in isolation and are surrounded and interact with many non-neuronal cells such as the various kinds of glial cells^3,4^. E.g. it has been well established that astrocyte-secreted factors promote synapse formation^5^ or that neuronal activity regulates oligodendrocyte^6^ and microglial proliferation^7^. While *in vivo* models have been invaluable approaches, it is hard to dissect cell-cell interactions in intact organisms. Moreover, many disease-relevant genes, including those associated with Alzheimer’s disease, are poorly conserved or even lack clear mouse orthologs^8^. Human cell models such as the ones derived from iPSCs would therefore be needed to study such species-specific mechanisms^9^. In order to study cell-cell interactions, it is thus critical to devise more complex human cell models that contain more than one brain cell type.

There are two principal approaches to obtain more complex and more physiological human neural cell models. One prominent approach is the generation of cerebral organoids which takes advantage of the remarkable self-organizational potential of iPSCs^10,11,12,13,14^. Such organoid systems produce three-dimensional cultures with similarity of embryonic tissue thus allowing the study of fundamental developmental processes^15,16^. However, challenges that organoid approaches face are high intra-and inter-line variability, slow maturation, late gliogenesis, and absence of non-neuroectodermal cell types such as microglia^17^. A second approach to increase the cellular complexity of human cell models is the combination of defined cell types into multi-lineage co-cultures. As differentiation protocols for various glial cells, such as astrocytes^18^, microglia^8^, and oligodendrocytes^19^, have improved over the past years, such neuron-glial co-cultures are becoming feasible^20,21,22^. For both the organoid and co-culture approach, iPSC-derived microglia have already been successfully incorporated.

However, microglial survival was dependent on the presence of cytokines, and their morphology and cell biological features did not resemble that of ramified microglia in the brain, suggesting a lack of proper trophic support from non-microglial cells present in these models.

Here, we have established a new tri-lineage co-culture system incorporating neurons, astrocytes, and microglia that can be established in two and three dimensions featuring consistent reproducibility, functional properties of all three cell types, and microglia in an apparently more physiologically state than observed in previous approaches.

## RESULTS

### Establishment of a self-sustained and defined human tri-lineage co-culture system

The human brain consists of many different cell types including neurons, astrocytes, and microglia. However, co-culturing these 3 cell types which require different media and survival factors is not trivial. As we set out to accomplish the co-culture of neurons, astrocytes, and microglia we faced the need to optimize several variables including various differentiation protocols to generate the three cell types, time points and sequence of cell type combination, maturation stage of each cell type, and cell seeding density, and finally the media composition including nature and concentrations of cytokines (**Figure 1A**). Through a systematic, iterative approach we found that (i) iPS cell-derived microglia (iMG) progenitor cells can incorporate better into neuron-astrocyte cultures than more mature microglia, (ii) iMG do not survive when directly co-cultured with astrocyte progenitor cells (APCs; **Figure S1A**), whereas matured astrocytes, which have been cultured without EGF/FGF for one week, do support microglial survival and promote their ramification, (iii) when plated on matured neurons and astrocytes, iMG survive and ramify in common neural media without exogenous cytokines. These insights allowed us to establish a robust and optimized protocol to generate tri-lineage neural co-cultures grown as adherent cells on culture dishes (2D co-culture) (**Figure 1B**).

**Figure 1.**
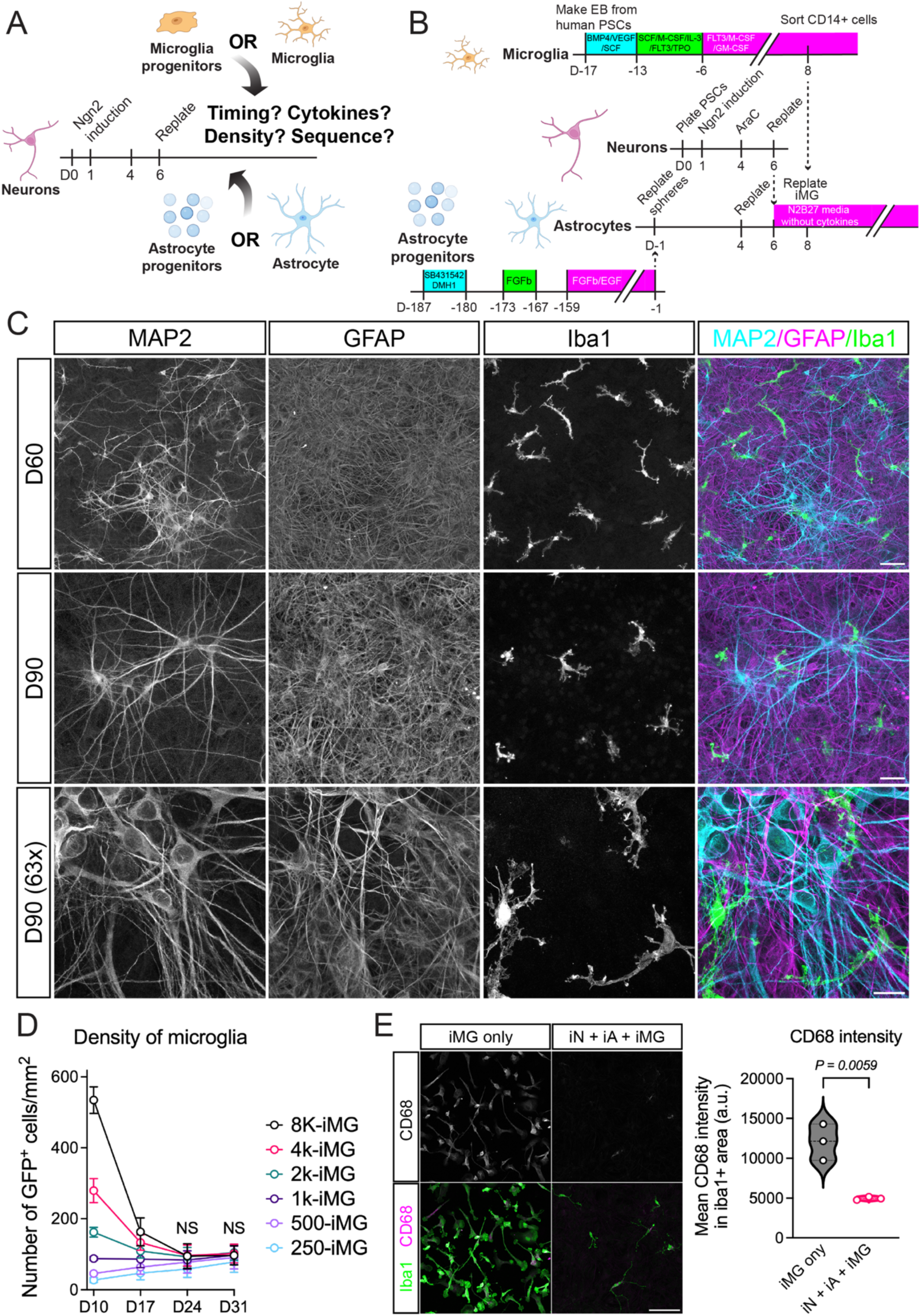
Establishment of a chemically defined and cytokine-free human neural tri-lineage co-culture system. (A) Illustration of the optimization process to develop neuron, astrocyte, and microglia co-cultures. (B) Outline of our optimized protocol to generate human pluripotent stem cell-derived tri-cultures. (C) Representative immunofluorescence images showing MAP2 (neurons), GFAP (astrocytes), and Iba1 (microglia) at indicated days after neuronal differention (see (B)). Scale bars, 50 μm (top and middle), 20 μm (bottom). (D) Quantification of microglial density over time following initial plating at varying cell numbers per well in a 384-well plate. *n* = 3 biological replicates, each with 3 technical replicates. Data are presented as mean ± SD. Statistical analysis was performed using two-way ANOVA followed by Tukey’s multiple comparisons test. NS, not significant. (E) Quantification of CD68 intensity within Iba1-positive areas in microglial mono-culture and tri-culture conditions. Microglia were cultured either alone (supplemented with 100 ng/ml IL-34 and 10 ng/ml M-CSF) or in tri-culture (without exogenous cytokines) for 14 days before fixation. *n* = 3 biological replicates. CD68 intensity in Iba1-positive areas was measured using ImageJ. Statistical significance was determined using an unpaired t-test.

Immunostaining for MAP2, GFAP, and Iba1 showed that all three cell types successfully incorporated into 2D cultures with complex neuronal, typical fibrous astrocyte and highly ramified microglial morphologies (**Figures 1C and S1B**). Next, we sought to evaluate the optimal seeding density of microglia. Surprisingly, we found that no matter how many microglia we seeded, in all cases the microglia density converged to about 10% of the total number of cells in the cultures after about two weeks (**Figures 1D, S1C, and S1D**). This observation suggests that the microglia density is intrinsically defined by the neuron-astrocyte component of the culture. We further found that microglia embedded in the tri-culture exhibited less expression of the lysosomal/phagocytosis marker CD68 compared to microglia in mono-culture suggesting the tri-culture environment provides a more physiological environment producing less activated and more homeostatic microglial cell states (**Figure 1E**).

### Microglia in the tri-lineage co-culture are dynamic, can phagocytose, and respond to inflammatory stimuli

Next, we wanted to explore the functional properties of microglia in our tri-culture system. First, microglia are known to survey the brain tissue by dynamically extending and retracting their processes *in vivo*^23^. Indeed, time-lapse imaging revealed that microglia in our tri-culture are highly dynamic as they continuously extend and retract their cell processes throughout the observation times resembling their behavior *in vivo* (**Figure 2A and Video S1**).

**Figure 2.**
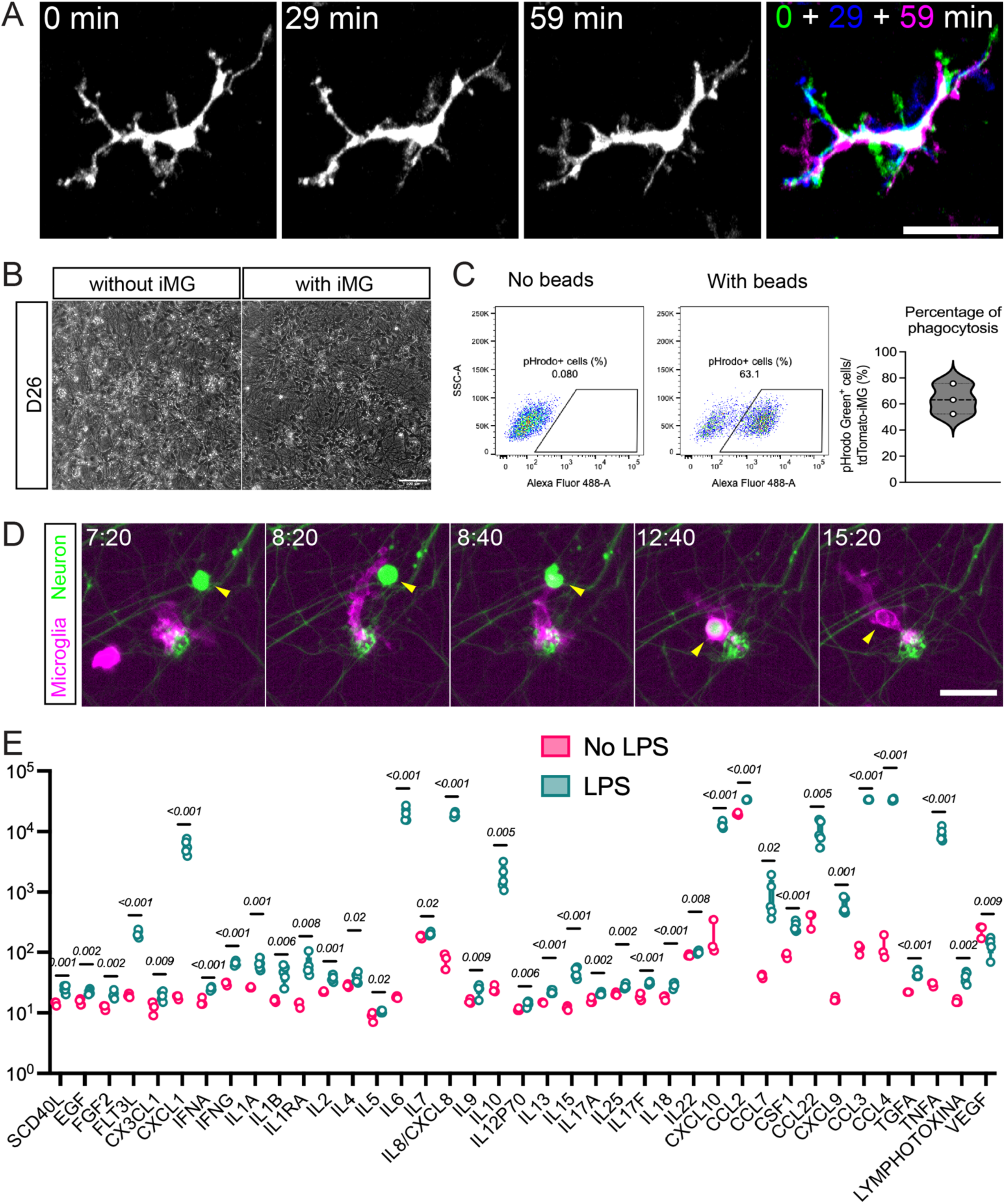
Functional characterization of microglia in the human tri-culture. (A) Time-lapse imaging of GFP-labeled microglia in DIV28 tri-culture revealed dynamic morphological changes in microglial processes. Scale bars, 50 μm. (B) Phase contrast images 26 days after differentiation. The presence of microglia in tri-cultures was associated with markedly less cellular debris. (C) Phagocytosis assay with pHrodo Green Zymosan A Bioparticles in tdTomato-labeled microglia at DIV29 tri-culture, measured by flow cytometry. *n* = 3 biological replicates. (D) Time-lapse imaging showed tdTomato-labeled microglia in DIV27 tri-culture phagocytosing GFP-labeled neurons. Microglia grasped the neuron at 8:40, enclosed the internalized neuron within its membrane by 12:40, and completed degradation of the neuron by 15:20. Numbers indicate hours:minutes. Scale bars, 100 μm. (E) Cytokine levels in the supernatant from DIV35 tri-culture were compared between LPS-treated and untreated conditions. *n* = 3–5 biological replicates. Statistical significance was determined using multiple unpaired t-tests.

Second, microglia are professional phagocytes. When comparing cultures with or without microglia, it became quite apparent that our tri-lineage cultures were almost completely devoid of any cellular debris suggesting an efficient clearance by microglial phagocytosis (**Figure 2B**). We confirmed their phagocytotic activity by using pHrodo-labeled Zymosan (a polysaccharide derived from yeast cell walls) and flow cytometry (**Figure 2C**) and observed microglia phagocytosing dying neurons in time lapse imaging (**Figure 2D and Video S2**).

Finally, a critical role of microglia is to respond to inflammatory stimuli and secrete immunoregulatory cytokines. To assess their inflammatory response, we added lipopolysaccharide (LPS) to microglia-containing tri-cultures and measured cytokines secreted into the media with or without the stimulus. Indeed, LPS induced the secretion of a large number of cytokines, including several well-known pro-inflammatory cytokines, such as TNFα, IL-6, IL-1β, and IFN-γ, regulatory cytokines such as IL-10, as well as chemokines including CXCL1, CXCL9, CXCL10, CCL3, CCL4, CCL7, and CCL22 (**Figures 2E and S2**).

These data demonstrate that the microglia in our tri-lineage, neural co-culture possess functional properties, including dynamic cell process motility, phagocytosis, and inflammation-induced cytokine release.

### Functional properties of neurons and astrocytes in the tri-lineage neural co-culture

We next sought to confirm that also neurons and astrocytes possess functional properties in our tri-lineage co-cultures. Immunofluorescence analysis demonstrated the presence of synapsin-positive puncta in close association with MAP2-positive neurites suggesting the formation of synapses and functional maturation of neurons in the culture (**Figure 3A**). Indeed, electrophysiological recordings demonstrated that neurons exhibited passive membrane properties characteristic for iPS derived neurons, including Na^+^ and K^+^ currents, membrane potential, input resistance, and capacitance (**Figures 3B and 3C**). Recordings in current-clamp mode revealed that the majority of patched cells spontaneously fired trains of action potentials (**Figure 3D**). Injections of increasing steps of current elicited typical, repetitive action potentials of characteristic shapes and electric parameters (**Figures 3E and 3F**). Finally, we were able to detect spontaneous postsynaptic events of characteristic asymmetric shapes when recorded in the voltage-clamp mode (**Figures 3G, 3H and 3I**).

**Figure 3:**
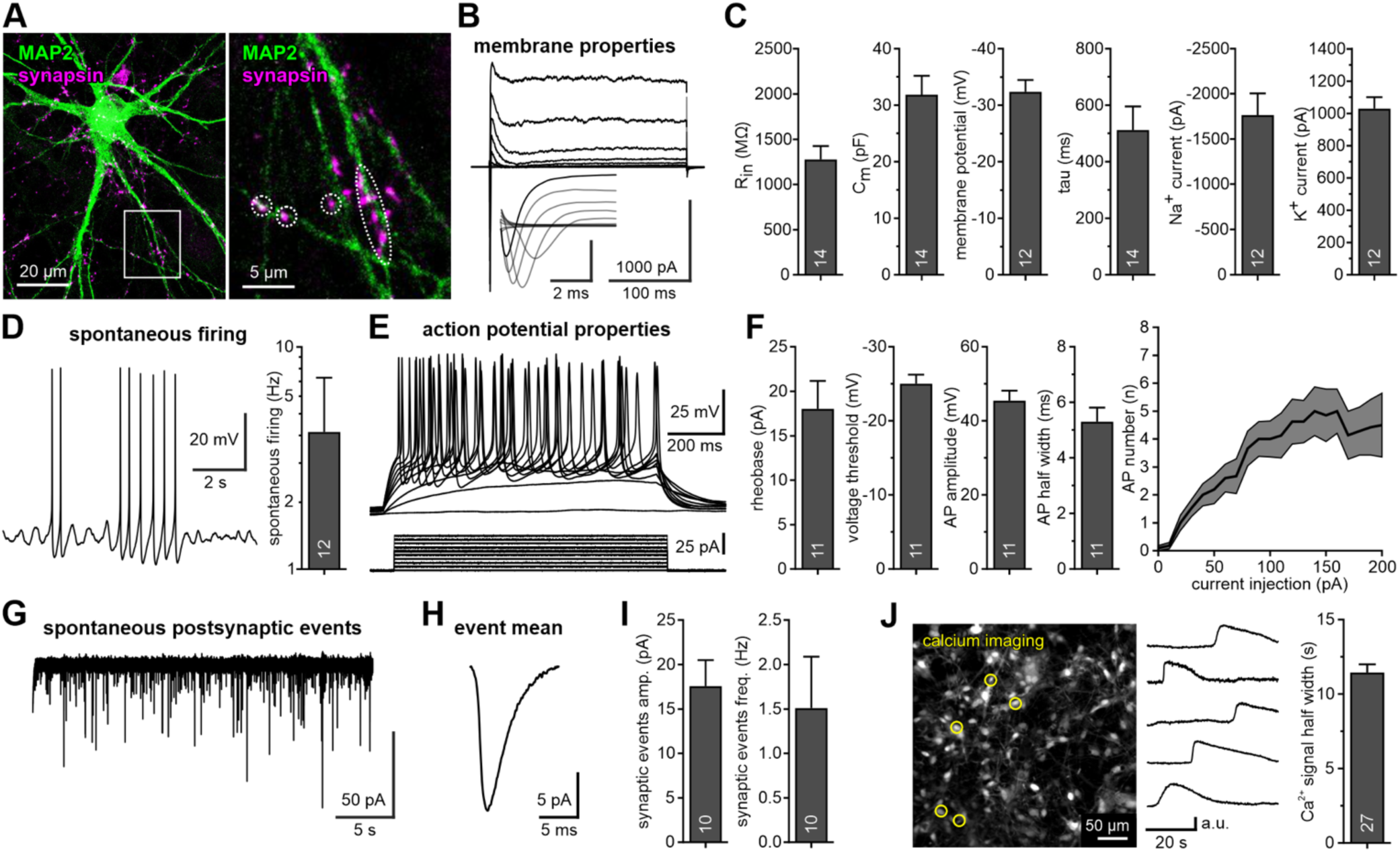
N**e**urons **and astrocytes in the tri-lineage co-culture exhibit functional properties** (A) Tri-cultures exhibit a mature neuronal network with synaptic junctions. Confocal detection of MAP2 (green) and synapsin (magenta) in a tri-culture 35 days after differentiation. The right panel represents a magnification of the region delineated by the white box. (B) Representative traces of voltage steps (-90 to +20 mV) analyzing voltage-gated sodium and potassium currents in patched neurons. (C) Assessment of passive membrane properties and peak currents from voltage-gated sodium and potassium channels. Rin=input resistance, Cm= membrane capacitance, tau= membrane time constant. (D) Sample trace of spontaneous action potentials (AP) and average spontaneous firing frequency. (E) Traces of action potential sequences elicited by increasing current injections (1 s duration, from 0 to 100 pA in 5 pA steps) of patched neurons. (F) Quantification of the action potential properties in tri-cultured neurons, including current and voltage thresholds, spike amplitudes and half-widths. Left panel shows number of action potentials evoked as well as quantification of action potentials evoked by increasing injections (as in panel E) plotted as I-F curve over a 1 s period. (G) Sample trace of spontaneous postsynaptic currents. (H) Mean of the postsynaptic events of cell depicted in G (note characteristic asymmetry). (I) Quantification of amplitude and frequency of spontaneous postsynaptic events. (J) Calcium imaging of astrocytes in tri-cultures using Fluo-8 AM dye. The left panel shows a standard deviation projection over a 90-second time-lapse, highlighting cells calcium transients. Traces illustrate fluorescence intensities of cells marked by yellow circles in the left panel. Bar plot on the right depicts mean half-widths of calcium trajectories from 27 individual cells All bar graphs in this figure represent mean ± SEM, with sample sizes indicated by white numerals within bars.

Synaptic maturation of neurons is known to require secreted factors derived from astrocytes^5^. Hence, the presence of functional synapses indicates that the astrocytes in the culture must have produced the proper pro-synaptic factors, an important astrocytic function. In addition, astrocytes are known to exhibit slow Ca^2+^ transients. To determine whether the astrocytes in our co-cultures possess this feature, we performed live Ca^2+^ imaging using a Ca^2+^-sensitive fluorescent dye. Indeed, we found many astrocytic cells in the visual field that exhibited characteristic slow changes of intracellular Ca^2+^ concentrations (**Figure 3J**).

### Neurons and glia approach transcriptional signatures of primary brain tissue

To assess and compare the transcriptional states of the cell populations of our tri-lineage co-culture with existing data sets, we performed single cell (sc) and single nucleus (sn) RNA-sequencing (seq) of 5-week-old cultures (about 3 weeks after microglia plating). We performed both approaches, since most human brain tissue has been analyzed by sn-RNA seq but transcript abundance is higher in sc-RNA seq. After initial quality control, cells were clustered and annotated using published marker genes. Both approaches revealed cell clusters that we could readily assign to the three cell types consisting of astroglia, neurons, and microglia (**Figures 4A and 4B**). We also detected an unexpected small cell population with transcriptional resemblance to fibroblastic stromal cells (**Figures 4A, 4B, 4C, and 4D**).

**Figure 4.**
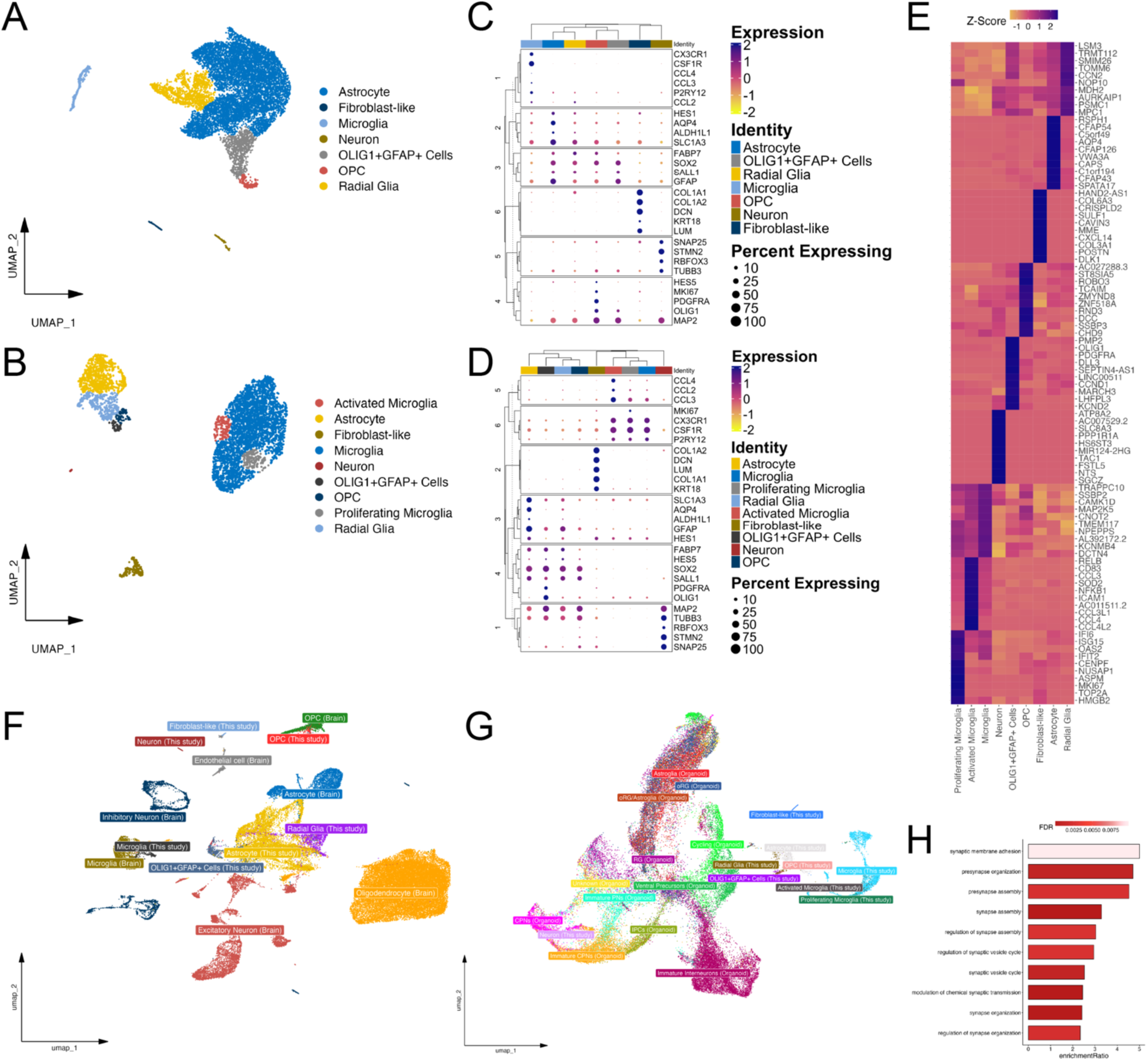
Single nucleus (sn) and single cell (sc) RNA-sequencing reveals the presence of various glial precursor cell types in the tri-lineage co-culture. (A, B) UMAP representation of (A) sn-and (B) sc-RNA seq with annotated cell populations. (C, D) Marker gene expression in each cell population of the (C) sn-and (D) sc-RNA seq data. Expression of each gene is scaled and color coded from yellow (low expression) to blue (high expression). The size of each dot represents the fraction of cells expressing the genes in each population. (E) Heatmap of the ten most specifically expressed genes in each population in the sc-RNA seq data. Expression of each gene is scaled and color coded from yellow (low expression) to blue (high expression). (F) UMAP representation of the tri-lineage co-culture sn-RNA seq sample integrated with *in vivo* brain samples. OPCs and Microglia of both datasets cluster together whereas the remaining cell populations of the tri-lineage co-culture cluster separately from their *in vivo* counterparts. (G) UMAP representation of the tri-lineage co-culture sc-RNA seq sample integrated with cerebral organoids. Neurons of both datasets cluster together whereas the remaining cell populations of the tri-lineage co-culture cluster separately from their *in vivo* counterparts. (H) Bar chart of neuron-specific enriched GO processes of the genes up-regulated in neurons of the tri-lineage co-culture compared to CPN brain organoid neurons. No neuron-specific processes were enriched for genes up-regulated in the CPN brain organoid neurons. The false discovery rate of each process is colored coded from low (dark red) to high (white).

*Cytokeratin 18* was one of the genes prominently expressed in these cells and we were able to detect clusters that stained positive with Cytokeratin 18 antibodies in our cultures. Lineage tracing experiments using tdTomato knock-in ESCs, revealed that this population is derived from our microglia differentiation, presumably a contamination of hematopoietic niche cells contained in the hemogenic embryoid body derivatives of the cultures (**Figure S3**). Purification of CD14^+^;CX3CR1^+^ microglia progenitor cells by FACS eliminated this stromal cell population in our cultures.

The astroglia in the culture were the most heterogeneous of the three cell types and, in fact, composed of multiple glial cell clusters (**Figures 4A and 4B**). In addition to mature astrocytes, the cluster analysis revealed the presence of more immature glial cells resembling radial glia, oligodendrocyte precursor cells, and a cluster containing progenitor cells characterized by prominent Olig1 and GFAP co-expression. Of note, we did not observe any more mature oligodendrocytic cells in neither transcriptional dataset and we failed to detect O4^+^ or MBP^+^ cells by immunofluorescence in the cultures.

Microglia represented a clearly distinct cluster. In the sc-RNA seq data a small proportion resembled proliferative, activated microglia (**Figures 4A and 4B**). The microglia expressed genes including the canonical markers CSF1R, P2RY12 and CX3CR1, and are characterized by the specific expression of TMEM117, NPEPPS and SSBP2 (**Figure 4E**). The proliferative microglia were characterized by expression of MKI67 and TOP2A as well as a characteristic high expression of IFI6, ISG15 and OAS2 (**Figure 4E**). In contrast, activated microglia expressed high levels of CCL2, CCL3 and CCL4 as well as NF-ΚB pathway genes such as NKFB1, RELB and ICAM1 (**Figure 4E**).

Even though live imaging and immunostaining demonstrated that the proportion of neurons is about 15% (**Figure S1D**), both sn-and sc-RNA seq experiments detected only a small handful of neurons. We presume that we lost most neurons during the single cell dissociation process which had to be rather harsh given the intricate network of mature neural tissue in these 5-week-old cultures. Neurons are particularly fragile presumably due to their complex and long neurites. Nevertheless, there were enough cells to obtain a sufficient gene expression coverage of the cluster. As expected, we found prominent expression of canonical marker genes such as RBFOX3, STMN2 andTUBB3 in the neuronal cluster (**Figures 4C and 4D**).

Moreover, we detected the expression of neuropeptide encoding genes TAC1 and NTS, the sodium/calcium exchanger SLC8A3 and MIR124-2HG, which promotes neuronal differentiation and represses non-neuronal genes (**Figure 4E**).

With single cell resolution in hand, we then explored the possibility to map the transcriptional profile of the cell types from our tri-lineage culture on primary cells from the brain. Most published datasets from primary human brain tissue are from sn-RNA seq studies. We therefore used our matching single nucleus data set as comparison. Specifically, we obtained datasets from normal appearing prefrontal cortex samples from Multiple Sclerosis patients^24^ and integrated them with our sn-RNA seq sample. Two populations, the OPCs and the microglia from our tri-culture clustered remarkably closely together with their respective counterparts from the human brain (**Figure 4H**). This result indicates that those two cell types are in fact quite mature and may reside in a transcriptional state that closely resembles their physiological counterparts. Nevertheless, in direct comparisons we observed transcriptional differences between the cell types from our culture and their *in vivo* counterparts (**Supplementary Table S4.1**). Gene Ontology (GO) enrichment of the upregulated genes for each comparison showed that only seven processes were enriched in microglia *in vivo*, relating to cell morphology (cytoskeleton-related terms) and microglia-neuron interactions (axon development). (**Supplementary Table S4.2**). Conversely, two GO terms related to extracellular structure organization were enriched in cultured microglia. These results underscore the inferred functional similarity of microglia in our culture and their *in vivo* counterparts. Surprisingly, for OPCs we observed many enriched GO processes. *In vivo* OPCs are enriched in neuron-glia communication, readiness for differentiation and plasticity, physiological roles tied to behavior and brain homeostasis as well as preserved morphology and adhesion mechanisms. Cultured OPCs show high translational and RNA metabolic activity and an upregulation of generic early developmental or metabolic programs. Both our neurons and astroglia were not clustering together with any primary cell types but they were distributed among neuronal and astroglial clusters from the brain (**Figure 4H**). This result suggests that our neurons and astrocytes are still clearly distinct from their *in vivo* counterparts. We presume that both cell types are less mature, which is supported by GO enrichment of cultured cells in generic cellular machinery, the absence of specialized function and increased proliferation-related activity (**Supplementary Table S4.2**).

Next, we sought to compare our tri-culture with the various cell types found in cerebral organoids. We obtained a published sc-RNA seq dataset from human cerebral organoids^25^, and used the corresponding sc-RNA seq from our culture as comparison. This time, we observed that our neurons clustered closely with organoid-derived “cortical projection” but not other types of neurons (**Figure 4I**). This finding is consistent with our previous findings that our NGN2-neurons are excitatory and express dorsal forebrain markers^2^. The astrocytes from the two culture systems on the other hand showed little overlap and gene expression changes (**Supplementary Table S4.3**). Although the genes upregulated in organoid-derived neurons were enriched in 74 GO terms, none of them was directly related to neuron function (**Supplementary Table S4.4**). In contrast, 10 of 74 enriched processes in our neurons were specific to neuronal functionality (**Figure 4J**). This result indicates that the two approaches are complementary and produces cells along a different maturation trajectory.

### Dissection of cell-to-cell communication mechanisms between different brain cell types

One of the advantages of our defined co-culture system is its modularity and the possibility to genetically manipulate each individual cell type selectively allowing for investigating cell-to-cell communication mechanisms. One of the prominent features of our culture system is that no exogenous cytokines are needed, demonstrating that the three cell types support each other’s survival and maturation. Hence, the question arose what the identity of those putative secreted factors are that interact between cell types guaranteeing homeostatic maintenance of the co-culture.

In order to identify specific ligand-receptor pairs responsible for neuron, astrocyte, and microglia survival within the culture, we utilized a recently developed computational tool, InterCom^26^, to predict cell-cell communications. Unlike conventional tools like “cell chat” this algorithm first pairs ligand-receptor cell types using a manually curated list with high interaction validity and then analyzes the transcriptome of the receiving cell type to confirm that the downstream pathway is actually activated. Following computation, we prepared predicted interaction lists from both the single cell and the single nuclear sequencing datasets for neurons, astrocytes, and (homeostatic) microglia (**Figure 5A, 5B, and Supplementary Table S5.1**). In total, we detected 124 interactions in the sc-RNA seq and 153 interactions in the sn-RNA seq data. We then defined the overlapping pairs and listed the ligand-receptor pairs with predicted active downstream signaling for all nine possible cell type pairs (**Figure 5C**). Most predicted interactions occurred in microglia-receiver pairs, followed by neuron-receiver pairs; astrocytes had no common incoming signals (**Figure 5C**). Coalescing these data based on the receptor molecule identity revealed that only eight receptors were responsible for all these predicted interactions (**Figure 5C**). These receptors included integrins, TGF-beta family receptors, axon pathway signaling molecules, and receptor tyrosine kinases. Due to the complementary nature of our sn-and sc-RNA seq, we also interrogated in how many interactions each receptor participates in the individual datasets (**Figure 5D and 5E**). Ten receptors were expressed by multiple cell types in at least one of the datasets (BMPR1A, EPHB2, FGFR1, IGF1R, IL6ST, ITGB1, ITGAV, NOTCH1, ROBO1 and TGFBR2), twenty receptors were expressed in a single cell type but involved in multiple interactions (ALK, BMPR1A, BMPR2, CD36, EPHB1, FAS, FGFR1, IGF1R, IL1RAPL1, INSR, ITGA2B, ITGA4, ITGAV, ITGB5, LRP1, LRP6, TGFBR1, TLR2, TLR4 and TNFRSF1A), and ten receptors were involved in unique interactions (CSF1R, EPHB2, FZD1, FZD2, IL1R1, LPAR1, NTRK2, PTPRF, ROR2 and UNC5B).

**Figure 5.**
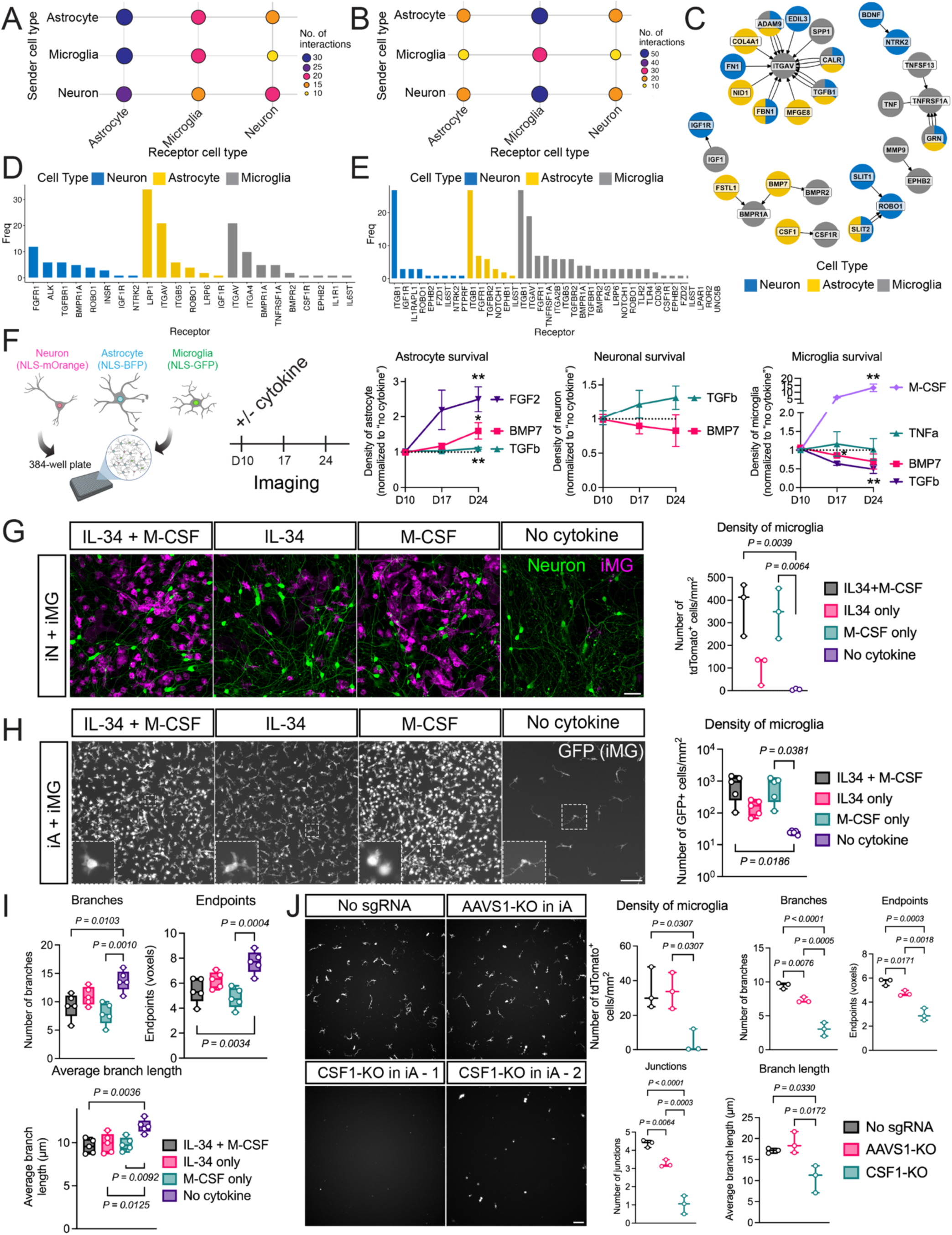
Cell-cell communication between neurons, astrocytes, and microglia in the human tri-culture. (A, B) Graphical representation of the number of predicted cell-cell interactions between astrocytes, neurons and microglia in the sn-(A) and sc-RNA seq data (B) of 5-week tri-lineage co-cultures. The amount of interactions between any two cell types are color coded from yellow (low) to blue (high) and scale proportionally with dot size. (C) Network representation of the common interactions identified in both the sn-and sc-RNA seq data. Node color corresponds to cell types. Multicolored nodes represent ligands that are expressed by multiple populations. (D, E) Bar chart of the number of interactions each receptor participates in per cell type in the sn-(D) and sc-RNA seq data (E). (F) Quantification of neuron, astrocyte, and microglia numbers under different cytokine conditions between days 10 and 24 of human tri-cultures. Cell numbers were normalized to wells without cytokine treatment and presented as mean ± SD. *n* = 4 biological replicates with 3 technical replicates each. Statistical significance was determined using two-way ANOVA with Dunnett’s multiple comparisons test. *, *p* < 0.05; **, *p* < 0.01. (G) (Left) Representative live image of a neuron–microglia bi-culture 10 days after microglia were added. Neurons were labeled with lentivirus expressing GFP, and microglia were differentiated from a human *CLYBL^CAG-tdTomato^* knock-in ESC line. Scale bars, 50 μm. (Right) tdTomato⁺ cells were quantified using ImageJ (n = 3 biological replicates). Statistical significance was assessed by one-way ANOVA followed by Tukey’s multiple comparisons test. (H) (Left) Representative live image of an astrocyte and microglia bi-culture 13 day after microglia were added. Microglia were differentiated from a *CLYBL^CAG-GFP^* knock-in human ESC line. Scale bars, 100 μm. (Right) The number of GFP^+^ cells was counted using ImageJ (n = 5 biological replicates). Statistical significance was determined using one-way ANOVA followed by Tukey’s multiple comparisons test. (I) Microglial morphology in (H) was quantified using the ImageJ plugin, MicrogliaMorphology. *n* = 5 biological replicates. Statistical significance was determined using one-way ANOVA followed by Tukey’s multiple comparisons test. (J) *CLYBL^CAG-tdTomato^* human microglia were co-cultured with unlabeled astrocytes, which were either not infected (no guide), or infected with LentiCRISPRv2 targeting the *AAVS1* or *CSF1* loci causing indel-mutations and loss-of-function mutations (see also Figure S4C). Images were taken 8 days after co-culture. Note near complete microglial absence or severe depletion and dysmorphic microglia upon astrocyte-restricted *CSF1* KO. Shown are one (no guide, AAVS1) and two (CSF1 KO) representative images of three biological replicates. Scale bars, 100 μm. tdTomato^+^ cells were quantified using ImageJ (n = 3 biological replicates). Microglial morphology was assessed using MicrogliaMorphology (n = 3 biological replicates). Statistical significance was determined using one-way ANOVA followed by Holm–Šídák’s multiple comparisons test.

To assess which of these pathways are involved in cell type-specific cell survival, we chose five readily available ligands known to activate many of the predicted receptors (FGF2, BMP7, TGFβ, TNFα, and M-CSF). Employing our previously developed live-imaging, cell type-specific counting assay (see also **Figure 1D, S1C and S1D**) we evaluated astrocyte, microglial, and neuronal survival over time. As predicted from the computational tool, FGF2, BMP7, and TGFβ significantly increased the density of astrocytes (**Figure 5F**). The addition of BMP7 or TGFβ also showed a trend to affect the density of neurons (**Figure 5F**). Microglia survival was significantly increased by M-CSF and decreased by BMP7 and TGFβ (**Figure 5F**), demonstrating the functional relevance of these three computationally predicted pathways in microglia.

The effect sizes of the functional ligand experiment were by far the strongest with M-CSF confirming the well-established critical role of CSF1R signaling for microglia. We next asked what the source of CSF1R stimulation is in our tri-culture. The computational tool predicted that microglial CSF1R is stimulated by M-CSF secreted from astrocytes (**Figure 5C**). We therefore first asked whether microglia survival depends on neurons or astrocytes. To this end, we plated microglia together with only neurons with or without CSF1R agonists. Quantification of microglia after 10 days of co-culture revealed that microglia survival is strictly dependent on the presence of cytokines (**Figure 5G**). Thus, neurons alone are incapable of supporting microglia. Next, we repeated the same experiment with astrocytes instead of neurons. This time microglia survived without cytokines in a similar density as in the tri-culture and also adopted much more ramified morphologies than in the presence of cytokines (**Figure 5H, 5I and S4A**). Therefore, astrocytes are clearly sufficient to support microglial survival and maturation. Next, we wondered whether the computational prediction is correct that it is astrocyte-derived M-CSF that contributes to this survival effect. We first validated using RNA-scope that M-CSF mRNA is indeed specifically detectable by hybridization in GFAP-expressing astrocytes (**Figure S4B**). To assess the functional role of astrocytic M-CSF, we then knocked out the *M-CSF (CSF1)* gene in astrocytes using two different CRISPR guides with a high frequency of frame-shift inducing indel mutations (**Figure S4C**). Indeed, unlike control guide treated cells, *CSF1*-mutant astrocytes were no longer able to support microglial survival and maturation (**Figure 5J**). Hence, we were able to confirm the prediction that astrocyte-derived M-CSF is responsible for microglia survival in our defined tri-culture.

### Establishment of a three-dimensional “tri-lineage brain microtissue”

Various three-dimensional culture models such as iPSC-derived organoids have been reported to support and maintain glia-neuron interactions^27,28^. However, incorporation of microglia into such systems have only been possible with exogenous cytokine support and resulting microglia often only adopt simple, atypical morphologies^22,29,30^. Given the strong microglial support of our co-culture system we wondered whether our system could also be applied in three dimensions. Rather than plating the three cell types sequentially to adhere to tissue culture dishes, we mixed and co-assembled them in U-shaped microwells previously coated with anti-adherent solution to support spontaneous aggregation (**Figure 6A**). In another iteration, we added microglia progenitors two to four days after assembly of neurons and astrocytes (**Figure 6A**). As a result, the cells reproducibly formed perfectly round, homogeneous microtissues and exhibited a similar appearance between wells within one differentiation run and between independent differentiation batches. The size of these microtissue spheres closely correlated with the number of cells plated. We chose a total cell number that yields spheres of a diameter of about 400 µm to ensure sufficient oxygenation throughout the structures (**Figure S5A**).

**Figure 6.**
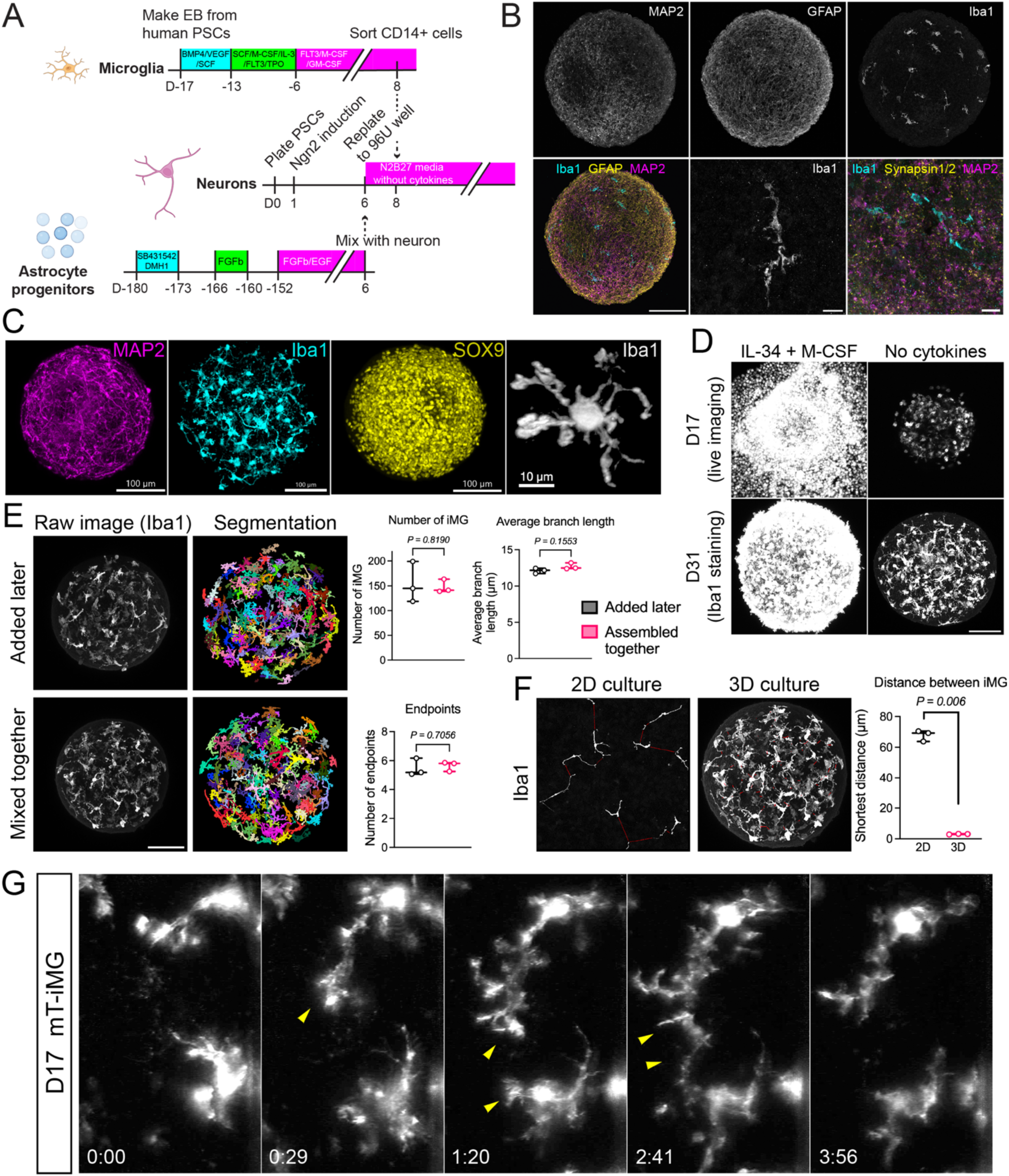
Establishment of tri-lineage brain microtissues. (A) Timeline of the assembly of human neurons, astrocytes, and microglia into microtissues. (B) Cryosections of brain microtissue at DIV33 stained with (top) MAP2, GFAP, and Iba1, and (bottom left) merged image. Scale bars, 100 μm. (Bottom middle) Higher magnification of an Iba1^+^ cell. (Bottom right) Co-staining of Iba1, Synapsin1/2, and MAP2 as indicated. Scale bars, 10 μm. (C) Whole-mount staining of cleared tissue showing MAP2 at DIV23 (left), Iba1 at DIV29 with microglia differentiated from an actin-tdTomato expressing line (second left), SOX9 (second right), and a representative image of an Iba1^+^ microglia at DIV29 (right). Images were processed and rendered using Imaris software. (D) Representative microglial images in brain microtissue with or without exogenous cytokines. Top: tdTomato-labeled microglia at DIV17. Bottom: Iba1-stained microglia at DIV31 after whole-mount staining and clearing. Scale bars, 100 μm. (E) Representative raw 3D volumes (left, Iba1+ in white) and corresponding cell segmentations used for quantification of tri-lineage brain microtissue with microglia either assembled at the same time with other cell types (middle-left bottom), or added later (middle-left top). Images were obtained 23 days after microglia were assembled. The timing of microglia addition does not significantly affect either the number of Iba1^+^ cells (middle-right, two-sided unpaired T-test, p=0.8191, T=0.255), or their key morphological characteristics (average branch length-top right, two-sided unpaired T-test, p=0.1553 (p=0.9314 after Bonferroni correction for non-independence), T=-1.763; number of endpoints - bottom right, two-sided unpaired T-test, p=0.7056 (p=1 after Bonferroni correction for non-independence), T=-0.414). (F) Representative raw 2D image (left) and 3D volume (middle) of 2D/3D tri-lineage cultures with Iba1^+^ microglia (white) and distances to the closest neighbor’s surface for each cell (red lines); the distance to their closest neighbor is significantly lower in 3D brain microtissue at DIV29 than in 2D tri-culture at DIV25 (right, two-sided unpaired T-test, p=0.0009666 (p=0.005799 after Bonferroni correction for multiple comparisons), T=-31.43). (G) Time-lapse imaging of membrane-tdTomato-labeled microglia in DIV17 brain microtissue. Two microglia extended their processes at 0:29 and 1:20, respectively. The two processes touched each other at 2:41 and then both retracted by 3:56. Numbers indicate hours:minutes.

Since the microtissues proved difficult to be penetrated by fluorescent light, we prepared cryo-sections and confirmed the presence of all three cell types by immunostaining with cell type specific markers (**Figure 6B**). We noticed that Iba1-positive microglia were evenly spaced throughout the entire sphere and even in these sectioned samples it became already apparent that the microglia displayed a high degree of ramification. To obtain a more holistic view of the three-dimensional organization of the microtissues, we subjected fixed samples to a clearing method^31,32^. Whole-mount staining with Iba1 antibodies of cleared microtissues revealed a morphological ramification of numerous processes even more enhanced than what we had observed in 2D cultures (**Figure 6C and Video S3**). Importantly, microglia showed a typical and homogenous tiling phenomenon throughout the microtissues. (**Figure 6C**). Sox9-positive astrocytes were homogenously distributed throughout the structures. As we had observed in the 2D culture, adding IL-34 and M-CSF greatly increased the microglial density and microglia lost their ramified morphology (**Figure 6D and S5B**). We did notice a substantial decrease in sphere diameter when microglia were added to the microtissues suggesting that microglia may clear debris from the structure (**Figure S5A**). Surprisingly, we also found that neuronal cell bodies migrated closer towards each other in the presence of microglia (**Figure S5F**).

We then compared the microglia incorporation pattern between the two different ways to add microglia to the microtissues, either by assembling all three cell types together or by adding microglia progenitors to existing neuron-astrocyte spheres. Computational segmentation and 3D reconstruction of the confocal Iba1 signal from cleared microtissues revealed that the number and morphological complexity of microglia were similar between the two conditions (**Figure 6E and S5E**).

Next, we sought to better compare the 2D and 3D incorporation pattern of microglia. Again, 3D reconstruction and nearest process distance measurements revealed that microglial processes are much closer to each other in the microtissues compared to 2D cultures (**Figure 6F and S5D**). This finding suggests the possibility that microglial processes may repel each other to accomplish cellular tiling as has been shown in the living brain^23,33^. Indeed, time lapse imaging of microglia derived from a tdTomato knock-in cell line incorporated into microtissues exhibited rather static cell bodies but highly dynamic processes which retract when encountering a process from a neighboring microglia (**Figure 6G and Video S4**).

### Human brain microtissues support formation of large α-synuclein inclusion bodies

Finally, we sought to confirm that our new model can be used to study known pathogenetic processes. The injection of α-synuclein preformed fibrils (PFF) into mouse brains or to primary cultures demonstrated the formation and spread of newly formed fibrils by recruiting endogenous monomeric α-synuclein which in turn leads to neurodegenerative features *in vivo*^34,35,36^. Defined culture models would be valuable to investigate mechanisms related to α-synuclein uptake, fibrilization, clearance, and spreading. However, most primary mouse neurons cannot be maintained for much longer than 3 weeks and human immature neurons only express low levels of α-synuclein. Since our human tri-cultures are long-lived and neurons become functionally mature we wanted to test whether α-synuclein pathology can be recapitulated in this system. To this end, we added various amounts of PFFs to a 3-week established 2D tri-culture and characterized the cultures with anti-phospho-synuclein antibodies (P-S129) which detects newly formed cellular fibrils but does not react with the synthetic PFFs (**Figure 7A**). Indeed, we observed large, round P-S129 positive inclusion bodies in MAP2^+^ neuronal somata in a dose-and time-dependent manner (**Figure 7B, 7C and S6A**). Since three-dimensional tissue organization could be relevant to recapitulate more aspects of neural cells, we next wondered whether our 3D microtissue spheres would also exhibit a similar pathology. Again, we treated established 3D microtissues containing all three cell types with different concentrations of PFFs and observed robust formation of large, P-S129-reactive, neuronal inclusion bodies in a dose-dependent manner (**Figure 7D, 7E, S6B and S6C**). Hence, our model will be useful to investigate key cellular mechanisms of α-synuclein pathology in a human cell context.

**Figure 7.**
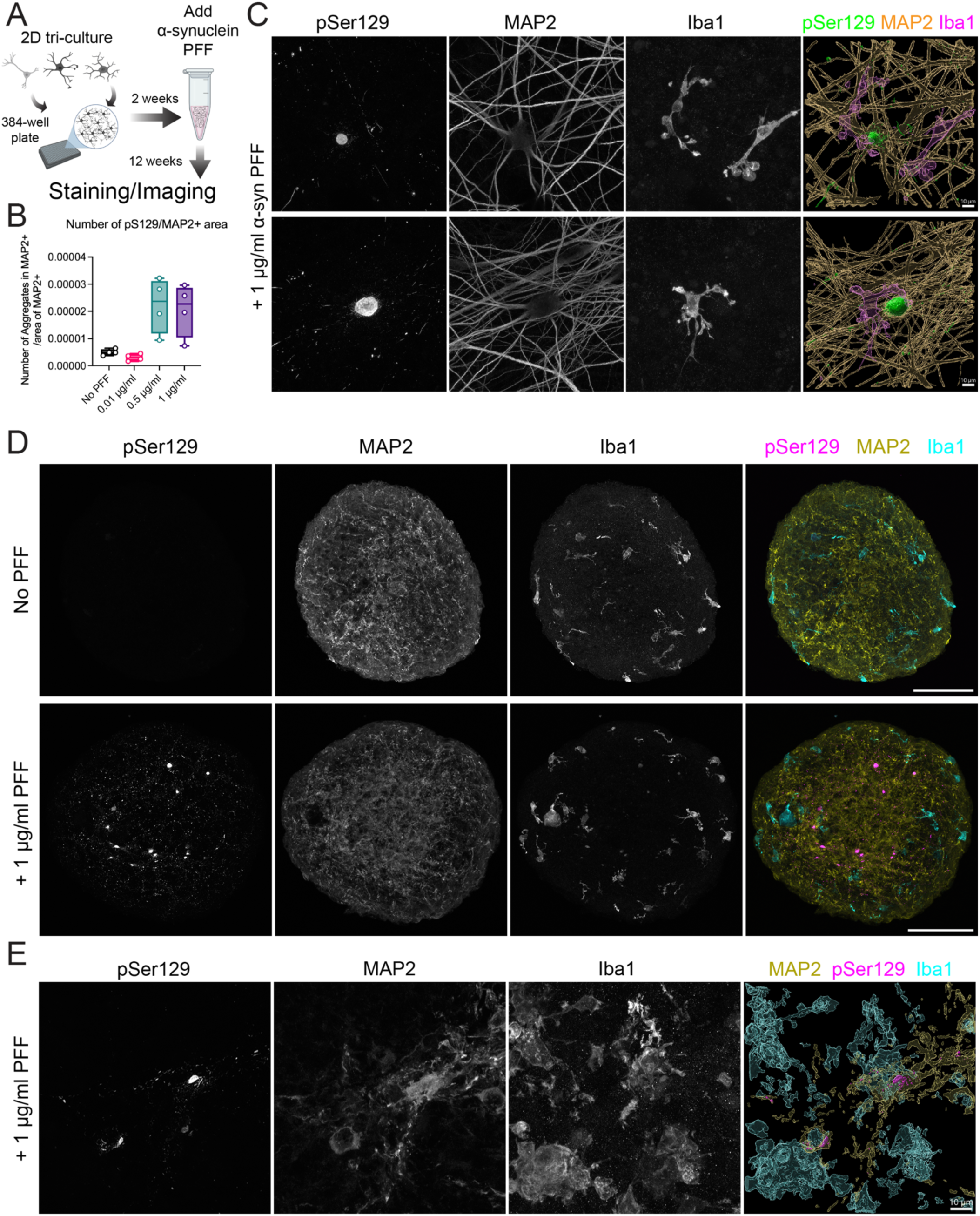
Neuronal α-synuclein inclusion body formation in human 2D and 3D tri-lineage brain microtissues following exposure to recombinant preformed fibrils (PFFs) (A) Experimental outline. Three cell types were plated in 384-well plate, and α-synuclein preformed fibrils (PFF) were added two weeks later. Cultures were stained and imaged 12 weeks after PFF addition. (B) The number of phospho–α-synuclein (pSer129)–positive dots localized in MAP2-positive areas was counted and normalized to the MAP2-positive area detected using ImageJ. Data are from one biological replicate with four technical replicates. (C) Representative images of 2D tri-lineage culture 12 weeks after PFF addition. (Left) Phosphorylated α-synuclein (pSer129), (second left) MAP2, (second right) Iba1. Merged images (right) were processed and rendered using Imaris software. (D) Representative images of 3D tri-lineage brain microtissue 4 weeks after PFF addition. The top panel shows the condition without PFF, and the bottom panel shows the sample treated with 1 μg/ml PFF. (Left) pSer129, (second left) MAP2, (second right) Iba1, and (right) merged images. Scale bars, 100 μm. (E) Representative images of 3D tri-lineage brain microtissue 4 weeks after PFF addition. (Left) pSer129, (second left) MAP2, (second right) Iba1. Merged images (right) were processed and rendered using Imaris software.

## DISCUSSION

In this paper, we developed novel tri-lineage co-culture systems consisting of human iPS cell-derived neurons, astrocytes, and microglia. With only slight procedural modifications the three cell types can form either two-dimensional tri-cultures or three-dimensional microtissues.

Several groups have already reported successful co-cultures of neurons, astrocytes, and microglia^20,22,37^. In these papers, the three cell types are generated separately, similar to this study, and then assembled in either 2D or 3D configurations. However, in these models microglia survival depended on addition of exogenous cytokines, e.g. IL-34 or M-CSF, well-known microglial survival factors that also affect functional states like activation and phagocytosis^38,39^. In our tri-culture systems microglia survived and exhibited mature phenotypes without any such exogenous cytokines. In fact, addition of exogenous cytokines caused pronounced cell proliferation and morphology changes. This finding suggests that the non-microglial cell types within the cultures provide the appropriate survival and maturation signals. Further investigation revealed that these signals are primarily derived from astrocytes in our culture system and that astrocyte-derived M-CSF is one of the necessary signals. For these reasons we speculate that the quality or maturation state of the astrocytes may be the most relevant difference between our and the other studies. Indeed, most studies generated astrocytes using rather rapid differentiation protocols sometimes including transcription factors to accelerated differentiation^40,41,42^. In contrast, the astrocyte progenitors we used here have been differentiated for at least 5 months and do not contain any progenitors with neuronal differentiation potential, which in our hands exist at earlier differentiation time points. Of note, one recent study did use very similar astrocytes and yet their microglia were cytokine-dependent^22^. This study used microglia differentiated with 6 transcription factors which were previously shown to be transcriptionally distinct from primary microglia, respond differently to some genetic perturbations, and thus may not have acquired a comprehensive microglial identity yet^43^.

One key observation of this paper is that the 3D context of our microtissues had a profound impact on microglial morphology and dynamic behavior. Three-dimensional cues may be required for full microglial maturation. Again, it is also possible that astrocyte maturation is a key contribution to microglial maturation in microtissues since astrocytes have been shown to exhibit a larger degree of maturation in 3D than 2D cultures^27^.

Surprisingly, cytokines were also required for the incorporation of microglia into cerebral organoids, which otherwise appear to authentically recapitulate many neurodevelopmental features^17,44,45,46^. Microglia during early neural development have a much lower density and different morphology^47,48^. Perhaps the source of microglial survival factors could be outside the developing neuroectoderm such as from the cerebrospinal fluid as proposed recently^49^. If mature astrocytes are in fact needed for proper microglial maturation, it would be not surprising that about 3-month-old cerebral organoids cannot support microglia as the development of mature astrocytes requires about 1 year of organoid culture^50^. Our 3D microtissues have additional helpful features compared to conventional organoids: We find minimal variability between microtissues; their size is experimentally controlled which avoids the central necrosis observed in long term organoid cultures impeding cell maturation and survival^51^; microtissues can be readily analyzed as whole mount preparations allowing the display of fully intact cells uncompromised by tissue sectioning^45,52^; the cell type composition is pre-defined and the different cell types can be generated from different stem cell lines allowing modular cell type composition, and finally the three cell types of the microtissues exhibit functional maturation within few weeks rather than many months after assembly.

Once proliferative ∼5-month astrocyte progenitor cells are obtained, the assembly of 2D or 3D tri-cultures is straight forward and accomplished in a few weeks. Since we combine immature neurons as early as day 6 after differentiation and microglia progenitors can be obtained as early as day 18 after differentiation, the rate-limiting step is the generation of astrocyte progenitors. For this reason, it is impractical to differentiate many different iPS cell lines into astrocyte progenitors. This is an obvious limitation of our specific method. However, astrocyte progenitor cells have a high proliferative capacity, can be cryopreserved, and can be used at least until ∼11 months after differentiation. A single differentiation batch can therefore be scaled exponentially which suffices for many experiments. Moreover, astrocyte progenitors can be readily infected with lentiviruses which allows genetic perturbation as shown here for lineage-tracing and loss-of-function applications (see e.g. **Figures 5F and J**). Genetic manipulation of astrocyte progenitors therefore represents a pragmatic alternative to their fresh differentiation from iPS cells.

For most versatility we explored both 2D and 3D tri-culture systems. Both versions have their respective strengths and weaknesses. 2D culture systems are better suited for live imaging, and live manipulation such as patch-clamp electrophysiology. 3D microtissues are easier to maintain long term cultures since media changes confer less mechanical stress on the cells than 2D cultured cells. They also provide cell surface for cell-to-cell interactions than 2D, which may contribute to the higher ramification of microglia and thus may enhance sensitivity to phenotypes that primarily driven by non-cell-autonomous interactions. Finally, whole-mount analysis of microtissues preserves the three-dimensional cell morphology. Therefore, our 2D and 3D cultures complement each other and serve as powerful tools for studying the cell biology of neurons, astrocytes, and microglia in a tri-lineage context.

Pathological protein aggregation is a common but still poorly understood phenomenon in neurodegenerative disease. In particular the mechanisms and cellular contributions of seeding and cell-to-cell spreading of pathological fibrils are unclear^53,54,55,56^. These mechanisms are hard to study in animal models such as rodents and additionally, many neurodegeneration-associated risk genes are poorly conserved between rodents and humans^57^. The *in vivo* spreading of pathological fibrils and subsequent neurodegeneration is particularly well established with a-synuclein^34^. We therefore sought to explore whether our defined human tri-cultures do recapitulate this pathological phenomenon. Remarkably, we observed robust formation of large, spheroid a-synuclein inclusion bodies in neuronal somata as well as elongated aggregates in neurites when seeded with pre-formed fibrils. The recapitulation of this critical pathological feature in our defined tri-lineage human cell models now opens the opportunity to investigate cellular mechanisms such as the contribution of microglia to aggregate formation.

In summary, we established defined human tri-lineage co-culture models featuring functional cellular properties, meaningful cell-cell interactions, and recapitulation of pathological protein aggregation. Our models will be useful the investigation of physiological and pathological processes that depend on non-cell autonomous mechanisms.

## LIMITATIONS

We capture three important cell types in our human cell model, but the human brain contains many more neuronal and glial subtypes along with endothelia and other vessel-associated cells. While all three cell types exhibit functional properties, compared to their *in vivo* counterparts, they are still less mature. Due to the extended differentiation times, the astrocytic component of our tri-cultures are impractical to obtain from different iPS cell lines. We recommend genetic modification at the astrocyte progenitor stage.

## ACKNOWLEDGMENTS

We would like to thank Catherine Crumpton from the FACS core at the Institute for Stem Cell Biology and Regenerative Medicine of Stanford University, Gordon Wang and David P Lenzi from Cell Sciences Imaging Facility of Stanford University for their assistance and advice. This work was funded by the NIH grants R01MH092931 and RF1AG048131, and grants from the Knight Initiative at the Wu Tsai Neuroscience Institute at Stanford University (KIG-105, KCG123). T.U. was supported by the Wu Tsai Neurosciences Institute Knight Initiative for Brain Resilience Scholar Award and Hereditary Disease Foundation Postdoctoral Fellowships. S.J. was supported by the Government of Canada through a New Frontiers in Research Transformation grant funded by the three federal research agencies CIHR, Natural Sciences and Engineering Research Council of Canada (NSERC) and Social Sciences and Humanities Research Council of Canada (NFRFT-2022-00327). M.R. is supported by Neuroscience Research Training Scholarship via the American Academy of Neurology (SPO 348983). W.L. was supported by Larry L. Hillblom Foundation (2025-A-214-FEL). T.C. was supported by a Neuroscience Research Training Grant (NIH T32MH020016) through the Stanford Neurosciences PhD program and a Stanford Interdisciplinary Graduate Fellowship affiliated with the Wu Tsai Neurosciences Institute.

## AUTHOR CONTRIBUTIONS

M.W. conceived the project. M.W., T.U., I.K., K.V., M.R., and Y.Y. designed the experiments. T.U., I.K., K.V., M.R., Y.Y., M.K., C.T., W.L., T.C., E.G., Z.S., S.Z., J.W., and M.A. performed the experiments. T.U., S.J., I.K., Y.Y., C.T., T.C., K.C., Z.S., A.V., A.-L.M.M., and M.J. developed experimental protocols, tools, and reagents or analyzed data. M.W., T.U., S.J., I.K., K.V., M.R., Y.Y., C.T., W.L., T.C., K.C., S.Z., J.M.S., T.W.C., M.A.R., H.A.L., M.C.B., T.C.S., A.D.S., and E.U. wrote or edited the manuscript.

## DECLARATION OF INTERESTS

M.W. is a co-founder of Neucyte Inc., a scientific advisor for bit.bio Ltd, co-founder and scientific advisor of Lytherian Therapeutics and Theseus Therapies. E.U. has financial interest in Synapticure. M.B. declares outside interest in DEM Biopharma and Stylus Medicine. T.C.S. is a co-founder of Neucyte Inc. and serves as an SAB member for BitBio Ltd. H.A.L. is the co-founder and chief scientific officer of ND BioSciences, Epalinges, Switzerland and received funding from Merck Serono, AC Immune, UCB, and AbbVie. TWC is co-founder and SAB member of Qinotto Inc., Teal Rise Inc., and Vero Biosciences.

---

**Figure S1.**
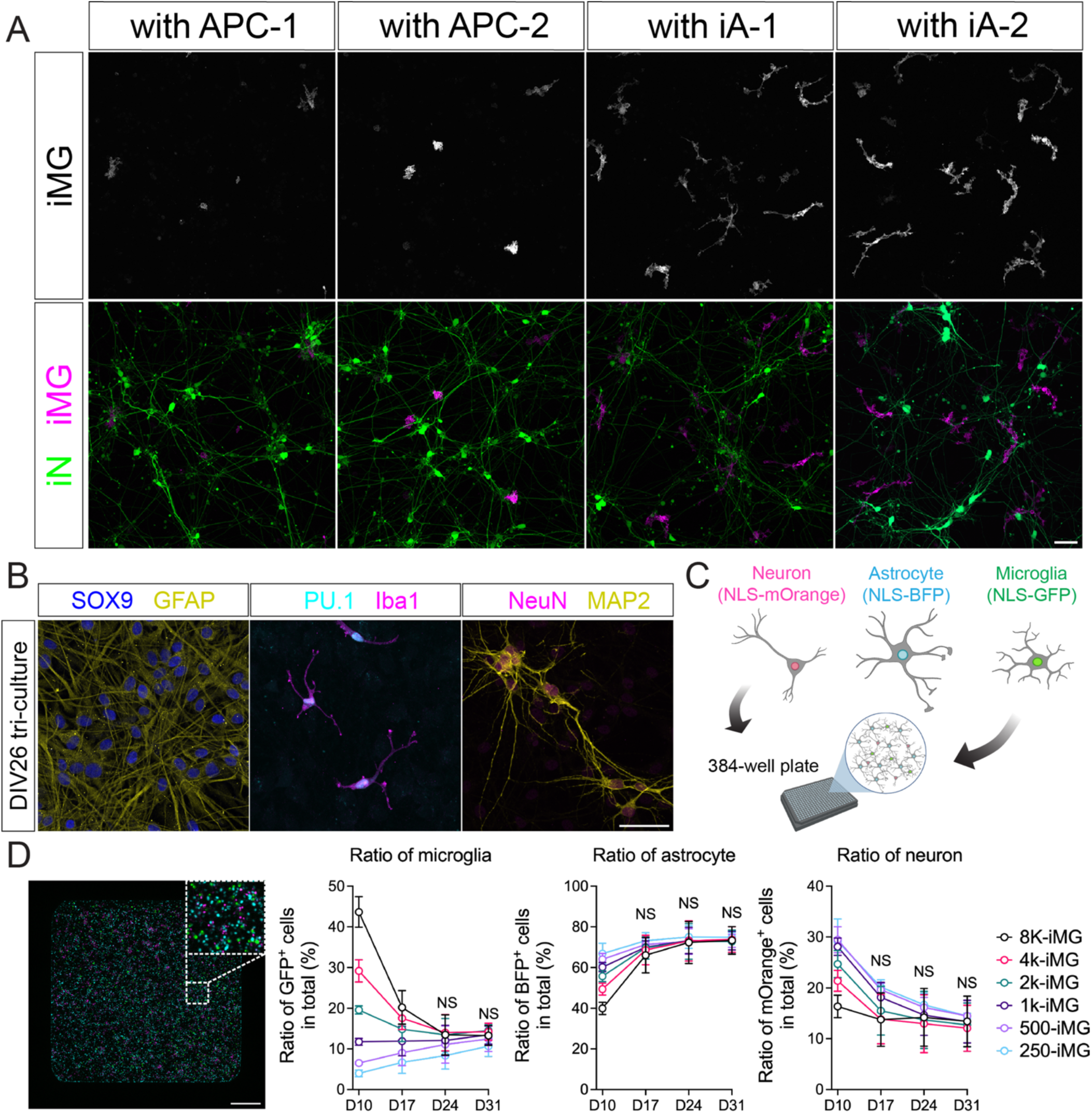
Optimization of the tri-culture system. (A) Representative live imaging of tdTomato-labeled microglia and GFP-labeled neurons co-cultured with either astrocyte progenitor cells (APC) or matured astrocytes (iA) at DIV18. Scale bars, 50 μm. (B) Representative immunostaining of 2D tri-lineage culture at DIV26. (Left) Co-staining with SOX9 and GFAP, (middle) PU.1 and Iba1, and (right) NeuN and MAP2. Scale bars, 50 μm. (C) Schematic illustration of the cell counting assay in a 384-well plate. NLS-mOrange– labeled neurons, NLS-BFP–labeled astrocytes, and NLS-GFP–labeled microglia were co-plated in each well. (D) (Left) Representative confocal image showing all three fluorescent markers. Scale bars, 500 μm. (Right) Quantification of microglia, astrocyte and neuronal ratio over time, percentage of total number. *n* = 3 biological replicates. Statistical analysis was performed using two-way ANOVA followed by Tukey’s multiple comparisons test. NS, not significant.

**Figure S2.**
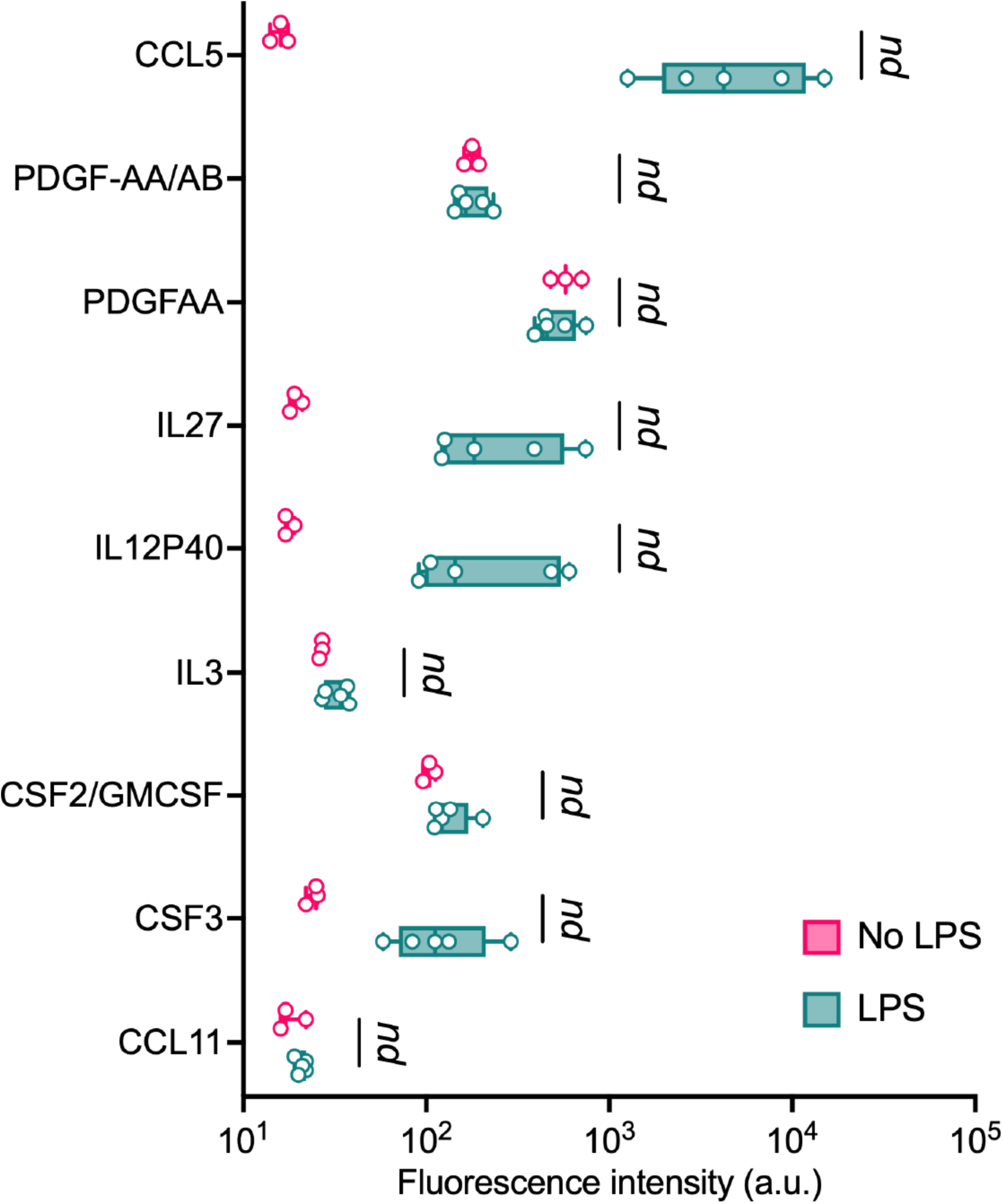
Cytokine secretion in response to LPS exposure. Additional cytokines measured in tri-culture with or without LPS treatment (not shown in Figure 2). *n* = 3–5 biological replicates. Statistical significance was determined using multiple unpaired t-tests.

**Figure S3.**
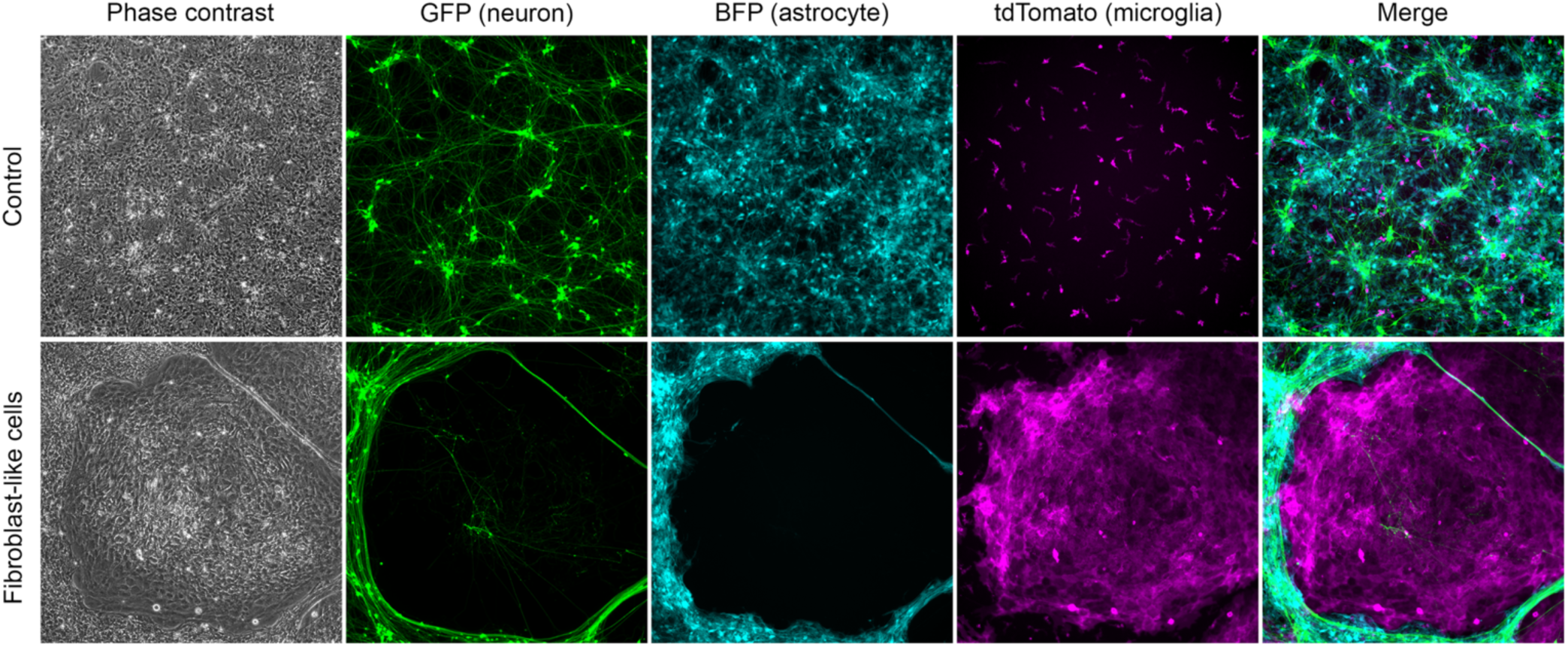
Stromal cell clusters in tri-lineage cultures are derived from the microglia differentiation protocol. Representative image of fibroblast-like cells derived from lineage-traced differentiation batches (GFP: neuronal differentiation, BFP: astrocyte differentiation, tdTomato: microglia progenitor differentiation) at day28 tri-culture. Microglia progenitors were enriched by MACS using CD14^+^ beads. The stromal cell-like clusters were not detectable when microglia were double sorted for CD14^+^/CX3CR1^+^ by FACS.

**Figure S4.**
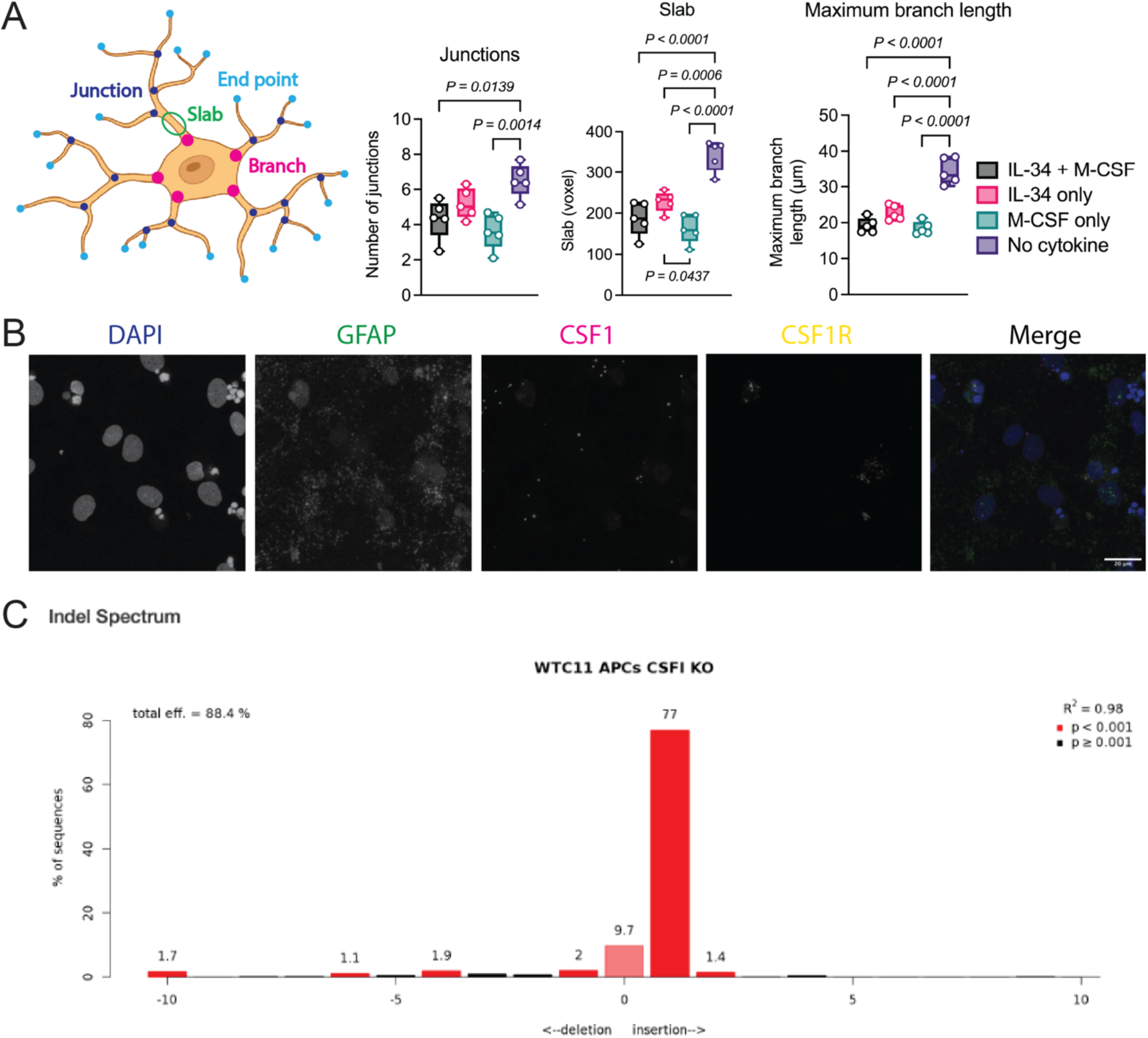
The relevance of M-CSF secreted from astrocytes. (A) (Left) schematic illustration for explaining where branch, junction, endpoint, and slab is. (Right) additional microglial morphological analysis in bi-culture of astrocyte and microglia. (B) Images show RNAscope for DAPI (nuclear marker) and probes for GFAP (astrocytes), CSF1 (expressed in astrocytes), CSF1R (expressed in microglia) in tri-culture 2D system. (C) WTC11 astrocytes infected with CSF1-targeting lentiCRISPR v2 and puromycin selected were analyzed with TIDE (https://tide.nki.nl), which detects the predominant mutations (i.e. deletions or insertions) as well as their frequencies. To do so, approximately 500 bp of the sgRNA mediated target sequence were amplified from control cells as well as targeted cells. The DNA fragments were sequenced by standard Sanger sequencing and compared on the TIDE platform.

**Figure S5.**
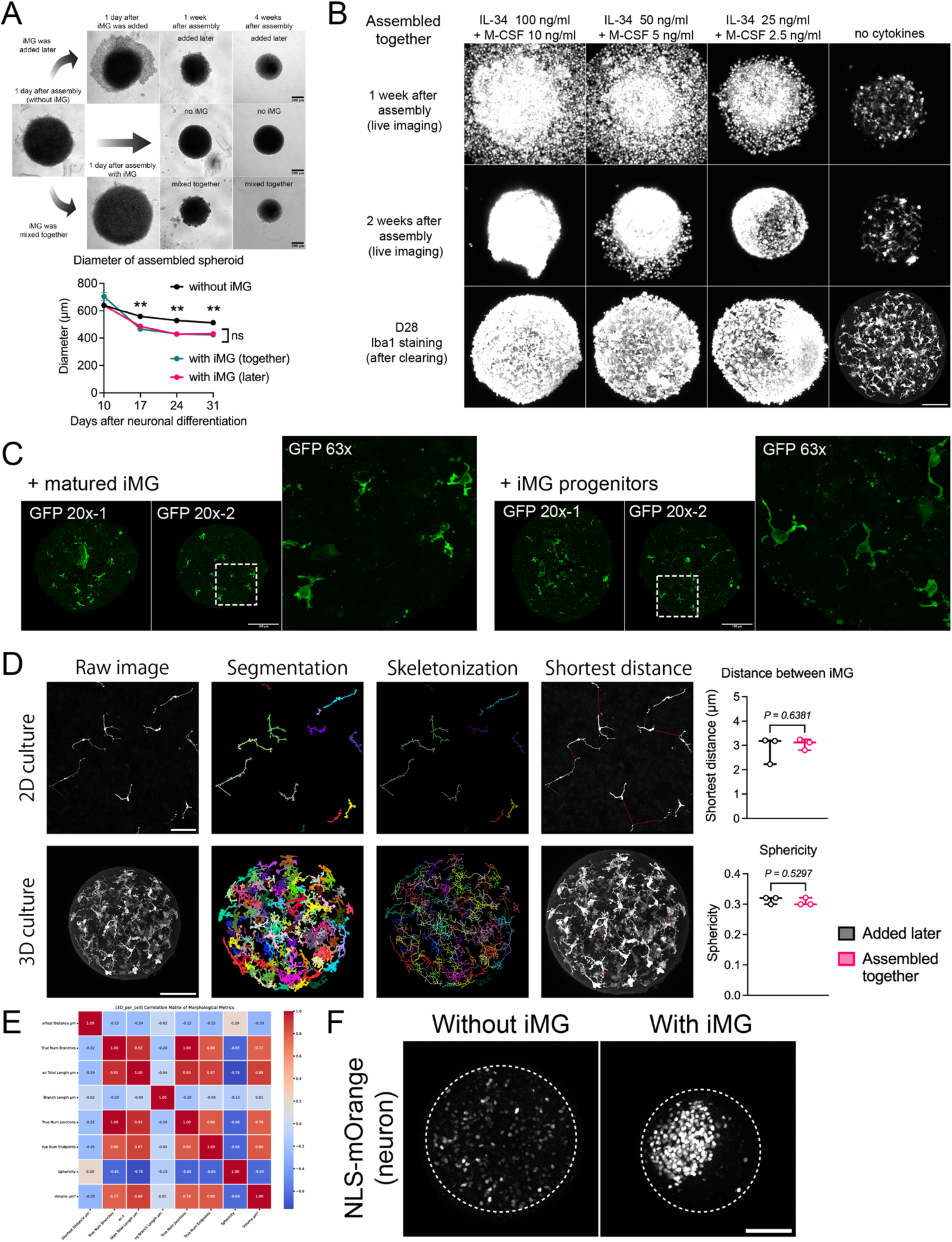
Optimization of brain microtissue formation. (A) Brain microtissue diameter measured weekly under two assembly conditions: microglia assembled together at DIV6 or added later at DIV9. *n* = 3 biological replicates. Statistical significance was determined using two-way ANOVA with Šídák’s multiple comparisons test. *, *p* < 0.05; **, *p* < 0.01. (B) Microglia in brain microtissues supplemented with different concentrations of IL-34 and M-CSF at 1, 2, and 3 weeks after co-culture. Top and middle: tdTomato-labeled microglia. Bottom: Iba1-stained microglia at DIV28 following whole-mount staining and clearing. Scale bars, 100 μm. (C) Immunostaining of DIV62 brain microtissue sections for GFP (to detect GFP-labeled microglia). Both matured microglia (left) and microglial progenitors (right) successfully incorporated into the brain microtissue. (D) Key output layers from the image analysis pipeline for a representative 2D image/3D volume (left). The timing of induced microglia addition into 3D tri-culture does not significantly affect either the distance between Iba1+ cells (top-right, two-sided unpaired T-test, p=0.6381 (p=1 after Bonferroni correction for multiple comparisons), T=-0.529), or their key sphericity (bottom right, two-sided unpaired T-test, p=0.5297 (p=1 after Bonferroni correction for non-independence), T=0.687). Scale bars, 100 μm. (E) A Pearson correlation matrix for the morphological heuristics extracted from the data used to identify the most orthogonal morphological properties for statistical analysis. (F) Representative images of NLS-mOrange–labeled neurons in DIV31 brain microtissue with or without microglia. Scale bars, 100 μm.

**Fig S6.**
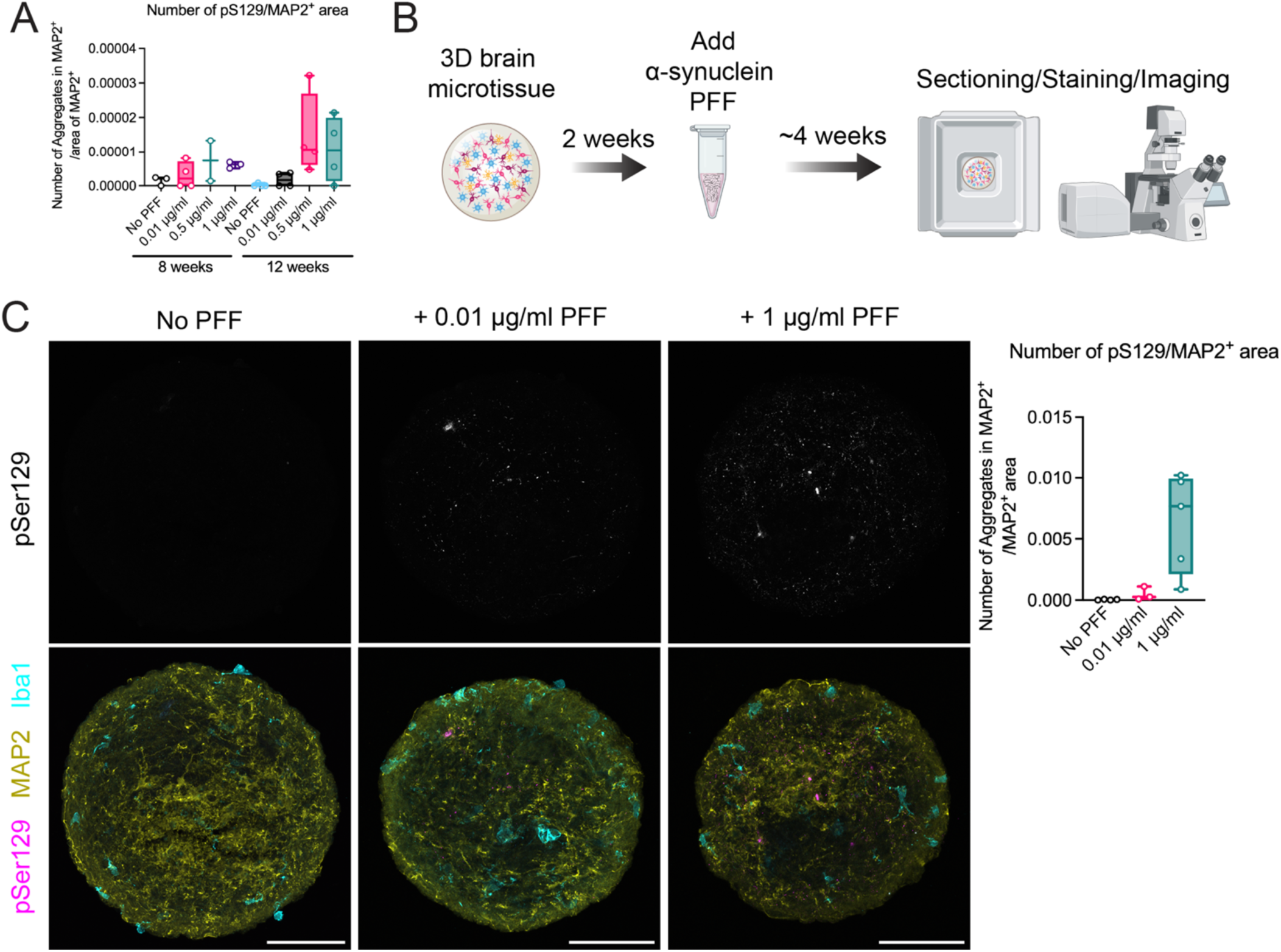
Optimization for establishing α-synuclein fibril inclusion body model in human tri-lineage culture by addition of α-synuclein PFFs. (A) Quantification of aggregates in MAP2-positive areas in 8-and 12-week-old 2D tri-lineage cultures. pSer129-positive dots localized in MAP2-positive areas were manually counted in a blinded manner and normalized to the MAP2-positive area detected using ImageJ. n = 1 biological replicate with 2 to 4 technical replicates. (B) Schematic of modeling α-synuclein aggregation in tri-lineage culture by addition of α-synuclein preformed fibrils (PFF). (C) 3D brain microtissues treated with α-synuclein PFF were sectioned and stained with antibodies against phosphorylated α-synuclein (pSer129) and MAP2 at DIV52. Representative images showing (top) pSer129 and (bottom) merged images with MAP2 and Iba1 at different concentrations of PFF. The number of pSer129-positive puncta localized in MAP2-positive areas was counted and normalized to the MAP2-positive area using ImageJ. Each dot represents one section analyzed from each of the 3–5 technical replicates. n = 1 biological replicate.

**Video S1. Time-lapse imaging of day 28 microglia in 2D tri-lineage culture, showing complex changes in ramified processes, related to Figure 2A Numbers indicate hours:minutes.**

**Video S2. Time-lapse imaging of day 27 microglia in 2D tri-lineage culture, showing a phagocytic event targeting an unhealthy neuron, related to Figure 2D Numbers indicate hours:minutes.**

**Video S3. Confocal Z-stack playback showing upward movement along the Z-axis of MAP2, Iba1, and SOX9 immunostaining in cleared brain microtissue, demonstrating successful staining throughout the entire sphere, related to Figure 6C** Brightness was adjusted in ImageJ to compensate for reduced laser penetration at deeper imaging planes.

**Video S4. Time-lapse imaging of day 17 microglia in 3D tri-lineage brain microtissue, demonstrating dynamic extension and retraction of processes, related to Figure 6G** Numbers indicate hours:minutes.

## METHOD DETAILS

### Pluripotent Stem Cell Culture

Male human embryonic stem cell (ESC) line H1 was obtained from WiCell Research Institute. The KOLF2.1J naïve iPSC line and KOLF2.1J containing TetO-Ngn2 and CAG-rtTA were obtained from Jackson Laboratory. The WTC11 iPSC line was obtained from the Gladstone Institutes. PSCs were cultured on Matrigel-coated plates in StemFlex medium.

### Generation of Ngn2-Induced Neurons

PSCs were differentiated with slight modifications to a previously described protocol^2^. KOLF2.1J-Ngn2 cells were plated as dissociated cells in a 6-well plate (1–2 × 10⁵ cells) on day 0. H1 cells without the Ngn2 cassette were plated with lentiviruses containing expression constructs for Ngn2-T2A-puromycin and rtTA.

On day 1-3, the culture medium was replaced with N3 medium (composed of DMEM/F12 [Thermo Fisher, #11320033], N2 supplement [Thermo Fisher, #17502048], and MEM Non-Essential Amino Acids [Thermo Fisher, #11140050], supplemented with insulin [Sigma, #I6634]) containing doxycycline (2 μg/mL, Sigma, #D9891) to induce Ngn2 expression.

On days 4 and 5, AraC (4 μM, Sigma, #C1768) was added to eliminate dividing cells. On day 6, neurons were dissociated and assembled with astrocytes as described below, in DMEM/F12 + GlutaMAX (Thermo Fisher, #10565018) supplemented with 0.5× N2 and 0.5× B27 without vitamin A (Thermo Fisher, #12587010), hereafter referred to as N2B27 medium.

### Human iPSC-derived astrocyte induction

Cortical-like astrocytes were generated from the human iPSC line WTC11 as previously described^58^. Briefly, on day 0 of differentiation, iPSCs were dissociated into small aggregates and transferred to untreated tissue culture flasks with the SMAD inhibitors SB431542 (Stemcell Tech, #72234) and DMH1 (Tocris, #73634). On day 7, when embryoid bodies began to show rosette clusters, the spheroids were transferred to Matrigel-coated tissue culture plates without SMAD inhibitors. On day 14, rosette clusters were mechanically removed and transferred to tissue culture flasks containing FGFb (Peprotech, #100-18B). On day 20, spheroids were triturated into a single-cell suspension and transferred to a new untreated tissue culture flask. From day 28 to day 180, spheroid aggregates were maintained in suspension with EGF (Peprotech, #100-15) and FGFb. The medium was changed every 4–5 days. Spheroid aggregates were triturated with Accutase every 7–10 days and transferred to new untreated tissue culture flasks.

### Differentiation for human PSC-derived microglia

PSCs were differentiated to microglia progenitors based on the previously described protocol^59^. 18 days after differentiation, supernatant was collected and CD14^+^ or CD14^+^/CX3CR1^+^ cells were sorted by FACS and used for further maturation in tri-culture.

### 2D bi-culture of PSC-derived neuron and microglia

The KOLF-Ngn2 line was infected with a lentivirus expressing EGFP and then differentiated into neurons. On day 6, 1.5 × 10⁴ neurons were replated into each well of a 96-well glass-bottom plate (Cellvis, #P96-1.5H-N) coated with poly-L-ornithine hydrobromide (Sigma, #P3655) and iMatrix-511 (Reprocell, #NP892-011). Two days later, 3 × 10³ microglial progenitors, differentiated from H1 cells stably expressing membrane-targeted tdTomato at the CLYBL locus, were added to each well. A half-medium change (N2B27 medium with appropriate cytokines) was performed every 2–3 days.

Images were acquired using an LSM710 confocal microscope, and the number of microglia was quantified using ImageJ.

### 2D bi-culture of PSC-derived astrocyte and microglia

Astrocyte precursor cells were dissociated, and 8 × 10⁵ cells in N2B27 medium supplemented with Y-27632 (but without EGF/FGF) were replated into a 6 cm dish coated with Matrigel.

One week later, astrocytes were detached using Accutase, and 1 × 10⁵ cells were replated into each well of a 24-well plate coated with Matrigel. Two days later, 2 × 10⁴ microglial progenitors, differentiated from H1 cells stably expressing EGFP at the CLYBL locus, were plated on top of the astrocytes.

A half-medium change (N2B27 medium with appropriate cytokines) was performed every 2–3 days. Images were acquired using a Leica DMi8 epifluorescence microscope, and the number of microglia was quantified using ImageJ. Microglial morphology was analyzed using the MicrogliaMorphology^60^ pipeline.

### Generation of 2D tri-culture

On day-3, astrocyte progenitors are dissociated as described above and plated at 2 × 10⁶ cells per 10 cm dish in N2B27 medium with Y-27632, but without EGF/FGF. The next day, Y-27632 is removed, and a complete medium change is performed every other day. On Day 4 (approximately one week after the initial dissociation), the culture is dissociated using Accutase and replated in N2B27 medium with Y-27632 at 5 × 10^4^-1 x 10^5^ cells per well of a 24-well plate. On Day 6, Ngn2-neurons are replated on top of the astrocytes at 1 × 10⁵ neurons per well of a 24-well plate. On Day 8, microglial progenitors sorted as described above are added to the neuron/astrocyte co-culture at 2 × 10⁴ cells per well of a 24-well plate. Half-medium changes are performed every 2–3 days.

### Generation of 3D tri-culture

The assembly of Ngn2-neurons and astrocytes is described in a previous paper^27^. Briefly, non– tissue culture-treated 96-well round-bottom plates (Corning, #3788) are coated with Anti-Adherence Rinsing Solution (StemCell Technologies, #07010). Astrocyte progenitor cells and Ngn2-neurons are dissociated on Day 6 after neuronal induction, then mixed at a 1:1 ratio (2 × 10⁴ cells each in 200 µL of N2B27 medium with Y-27632 per well of a 96-well U-bottom plate). The plate is centrifuged at 300 × g for 3 minutes. Half-medium changes are performed every 2–3 days.

Microglial progenitors (4 × 10³ cells), sorted as described above, can either be mixed together with neurons and astrocytes at the time of assembly or added a few days later, as shown in Fig. 6.

### Immunostaining for 2D tri-culture

Coverslips or entire tissue culture plates were fixed with 4% paraformaldehyde (PFA) for 10 minutes at room temperature (RT). After washing with DPBS, cells were permeabilized with 0.2% Triton X-100 in DPBS for 8 minutes, followed by incubation with blocking buffer (5% BSA in DPBS with 0.1% Triton X-100) for 1 hour at RT. Cells were then incubated with primary antibodies diluted in blocking buffer overnight at 4°C. The next day, after DPBS washes, cells were incubated with secondary antibodies diluted in blocking buffer for 1 hour at RT. Finally, coverslips were washed with DPBS and mounted using DAPI Fluoromount-G^®^ (SouthernBiotech, #0100-20). Images were acquired using a ZEISS LSM980 confocal microscope.

### Cell counting assay in 384-well plate

The KOLF-Ngn2 line was labeled with a lentivirus carrying CAG-NLS-mOrange and then differentiated into neurons. Astrocyte precursor cells were infected with a lentivirus carrying CAG-NLS-mTagBFP2 upon dissociation, and subsequently plated in 6 cm or 10 cm dishes to form a monolayer. A piggyBac vector expressing CAG-NLS-EGFP was used to generate a polyclonal EGFP-expressing microglial line. The three cell types were mixed in a 384-well plate (7.5 × 10³ neurons and astrocytes, and 1.5 × 10³ microglia per well), as described above. Two days after microglia was added, confocal images were acquired and then captured weekly. The number of cells expressing each fluorophore was quantified using ImageJ.

### Time-lapse imaging in 2D tri-culture (Figure 2A)

Day 28 tri-cultures were imaged using a ZEISS LSM980 laser scanning confocal microscope equipped with Airyscan 2, a motorized XY stage, Definite Focus 2, and environmental control.

16.5µm depth Z-stacks of fluorescent microglia were captured with a 10x objective over the course of 1 hour with 1-minute intervals. Airyscan processing and maximum intensity projections were processed using Zen 3.3 software. Brightness/contrast was adjusted and 3 frames/second video files were created using ImageJ.

### Phagocytosis assay

The 2D tri-culture was prepared as described above, using 5 × 10⁴ astrocytes, 1 × 10⁵ Ngn2-neurons, and 6.7 × 10³ microglial progenitors per well in a 24-well plate. On day 29 after neuronal induction, tri-cultures were incubated with 0.05 mg/mL pHrodo™ Green Zymosan BioParticles™ Conjugate (Invitrogen, P35365) in N2B27 medium at 37 °C for 1 hour, following the manufacturer’s instructions. After incubation, the cells were washed once with PBS and detached using Accutase. The cell pellet was collected by centrifugation, resuspended in N2B27 medium with Y-27632, and gently dissociated by pipetting. After centrifugation to remove the medium, cells were resuspended in flow cytometry staining buffer (Invitrogen, 00-4222-26) containing Y-27632 and analyzed for phagocytic activity using a BD FACSAria II flow cytometer. Data were analyzed using FlowJo software.

### Time-lapse imaging for phagocytosis (Figure 2D)

EGFP-labeled iN, unlabeled astrocytes, and membrane tdTomato-labeled microglia were seeded on 35 mm glass-bottom dishes (MATTEK, P35G-1.5-20-C). On day 27, the cells were transferred to a Zeiss Axio Observer Z1 microscope equipped with an incubation chamber and imaged for 67 hours at 37°C and 5% CO2. Brightfield and fluorescence images were collected at multiple positions every 20 minutes using an automated stage controlled by the Micro-Manager software. We used a Zyla 5.5 sCMOS camera and an A-plan 10x/0.25NA Ph1 objective. Movies were created using ImageJ.

### Measurement of cytokine

The 2D tri-culture was prepared as described above. On day 34 after neuronal induction, a complete medium change was performed, and fresh N2B27 medium with or without LPS (1 µg/mL, Sigma, LPS25; E. coli O111:B4) was added. After 24 hours, the supernatant was collected and centrifuged at 1,400 rpm for 10 minutes at 4 °C. The supernatant was then stored at –80 °C. Cytokine levels were measured using the Luminex assay at the Human Immune Monitoring Center, Stanford University School of Medicine.

### Electrophysiology in tri-culture

Electrophysiology was performed on neuron in tri-cultures 35 days after neuronal induction. Cells were visualized with a 40x water immersion objective (NA 0.8) using infrared oblique illumination optics (Olympus BX51WI). Neuronal identity was confirmed using an intrinsic fluorophore expression. Recordings were filtered at 10 kHz and sampled at 50 kHz using an Axon MultiClamp 700B amplifier with a Digidata 1440 digitizer using the Clampex 10.5 software (Molecular Devices, San Jose CA). Patch pipettes (borosilicate glass, OD 1.5 mm, ID 1.12 mm; TW150-4, WPI, Sarasota FL, USA) were pulled on a Narishige PC-10 puller (Narishige, Tokyo, Japan). Tip diameter was assessed electronically and ranged from 3 to 4.5 MΩ in the bath. Series resistances were compensated using the bridge circuity of the amplifier. The liquid-junction potential created by solutions of different ion compositions was not corrected for but calculated to be +14.5 mV. In current clamp, apart from the measurement of the resting potential, membrane potentials were adjusted by constant current application to values around-60 mV. Only cells with stable access resistances below 20 MOhm were used for analysis.

The internal solutions contained (in mM): 137 K-gluconate, 5 KCl, 10 HEPES, 0.2 EGTA, 10 Na-Phosphocreatine, 4 MgATP, 0.5 NaGTP, equilibrated to pH 7.3 with KOH. Membrane properties and ion currents were recorded in voltage clamp. Passive properties were captured using Clampex in the pCLAMP 10 Software Suite. Voltage-gated sodium and potassium currents were triggered by increasing 200 ms long voltage steps (-90 to +20 mV, 10 pA increments). Spontaneous synaptic activity was recorded for 5 minutes in the absence of any blockers. Action potentials were recorded in current clamp. Spontaneous action potentials were recorded for 1 minute at resting membrane potential. Firing characteristics and waveform properties were analyzed using increasing current steps (1000 ms length, 5pA increments up to 150 pA). Action potential waveforms were analyzed in the first action potential that fired 50ms after current injection onset. Rise and decay times were calculated from 20-80% of the current amplitude maximum, respectively. Halfwidths were calculated from 50% to 50% of the maximum amplitude.

### Single nuclear-and single-cell RNAseq

For single-nuclear sample collection, we adapted the protocol described by 10x Genomics://cdn.10xgenomics.com/image/upload/v1660261285/support-documents/CG000124_Demonstrated_Protocol_Nuclei_isolation_RevF.pdf]. Briefly, we prepared lysis buffer for single cell suspension (10mM Tris-HCL pH7.4, 10mM NaCl, 3mM MgCl2, and 0.025% NP-40). We prepared nuclei wash and resuspension buffer (1% BSA, 0.2U/μl RNase Inhibitor, in 1x PBS). To harvest nuclei, we removed culture media, washed once with 1x PBS, and added ice-cold lysis buffer (scaled up to accommodate larger cell quantities) and incubated the samples for 1 minute. Next, we collected the samples and added nuclei wash and resuspension buffer and mixed with gentle pipetting. The samples were spun down at 500g at 4°C and the supernatant was removed and discarded. Next, 1mL of ice-cold nuclei wash and resuspension buffer was added and mixed with gentle pipetting. The samples were spun down again at 500g and the supernatant was discarded. As a final step, the samples were resuspended in nuclei wash and resuspension buffer at a final concentration of 1,000 nuclei/μl (determined via manual counting with Trypan Blue staining) and used immediately in the 10x Genomics Single Cell Protocol. Leftover samples were flash-frozen in liquid nitrogen.

For single-cell sample collection, we followed the 10x Genomics cell preparation protocol (https://cdn.10xgenomics.com/image/upload/v1686678481/support-documents/CG00053_Handbook_CellPreparation_SingleCellProtocols_Rev_D.pdf). Briefly, we removed media from the samples, washed once with 1x PBS, and incubated with Accutase for 5 minutes at 37°C. The samples were checked at 2.5 minutes to ensure dissociation was proceeding as expected. 2mL of tri-culture Media was gently added to the samples, which were transferred to a 15mL conical tube and spun down at 20g for 5 minutes at room temperature in a swinging bucket centrifuge. The supernatant was discarded and the pellet resuspended gently in 1x PBS before being strained through the MACS SmartStrainer (30μm), run through a tabletop cell sorter to confirm the absence of doublets and clumps, and adding the resulting cells to the 10x Single Cell workflow.

scRNA-seq was performed using the Chromium Single Cell 3′ GEM, Library & Gel Bead Kit v3, 16 rxns (10x Genomics, PN-1000075) according to the manufacturer’s instructions and targeting a recovery of 10,000 cells per dataset. The libraries were constructed as instructed in the manufacturer’s protocol and sequenced by Novogene using NovaSeq PE150. Sequencing reads were processed with STARsolo (v.2.7.3a) using the human reference genome.

From the filtered barcode and count matrices, downstream analysis was performed using R (v.4.0.3). Quality control, filtering, data clustering, visualization and differential expression analysis were performed using the Seurat (v.4.0.3) and DoubletFinder (v.2.0) R packages. Datasets were processed following Seurat standard integration protocol according to the tutorial instructions. Genes expressed in less than 3 cells and cells with fewer than 500 features, less than 2,000 transcripts and more than 20% reads mapping to mitochondrial genes as well as cells identified as doublets by DoubletFinder were removed. PCA was performed for dimensionality reduction and the first 30 components were used for UMAP embedding and clustering.

### Ligand-receptor pair network

For reconstructing cell–cell communication networks, we used the single-cell-based method, InterCom, as described previously^26^. The main workflow of InterCom involves the following steps. First, for each cell type, transcription factors (TFs) with conserved expression across a subset of cells are identified. To this end, expression data is binarized: TFs with at least one count are assigned a value of 1, while non-expressed TFs are assigned −1. The top five percent of TFs with the most conserved expression are selected as preserved TFs.

Next, InterCom identifies intermediate signaling molecules that induce these preserved TFs using a Markov Chain model of intracellular signaling. These intermediates are linked to the initial TFs in the signaling cascade, and their transcriptional compatibility is evaluated. An intermediate molecule is considered compatible if its expression, together with the expression of downstream TF targets, aligns with the expected direction of the signaling pathway in a statistically significant number of cases (Hypergeometric test, p < 0.05).

Receptors are similarly connected to intermediate molecules if a compatible signaling path exists. Only receptors that are co-expressed with their interface TFs and downstream targets in at least 5% of cells are retained.

Finally, ligand–receptor interactions are established between cell types if the receptor has a verified downstream effect, the ligand is expressed in more than 5% of cells, and the interaction is included in the predefined interaction scaffold. An interaction score is calculated as the product of the average receptor expression and the average ligand expression across all cells expressing the respective receptor or ligand. The significance of interactions is determined by comparing scores across all potential cell–cell interactions within the scaffold, with those in the top decile considered significant.

### Generation of CSF1-KO astrocytes

For individual CRISPR/Cas9-mediated gene-disruption experiments, sgRNAs (sgAAVS1: GCCAGTAGCCAGCCCCGTCC, sgCSF1: GGCGAGCAGGAGTATCACCG) were cloned into lentiCRISPR v2 (Addgene #52961, deposited by Feng Zhang) as described previously^61^.

### RNAscope

RNAscope experiments were performed according to their protocol for Adherent Cells Cultured on Coverslips and the Multiplex fluorescent v2 Assay. In brief, cells cultured on coverslips were once rinsed with 1X PBS, followed by a 30 min fixation in 4% paraformaldehyde at room temperature (RT). We then performed dehydration by submerging the coverslips in 50% ethanol at RT for 1 min, 70% ethanol at RT for 1 min, and 100% ethanol at RT for 1 min. The 100% ethanol was replaced with fresh 100% ethanol. The coverslip was removed from ethanol and air dried to immobilize onto the glass slide by placing a drop of nail polish and then placing the upper edge of the coverslip cell-side up on the nail polish and allowed to air dry for 2-3 mins. A hydrophobic barrier was created around the coverslip using the Immedge™ hydrophobic barrier pen. The slides were rehydrated by placing them in 100% ethanol at RT for 1 min, then 70% ethanol at RT for 1 min, then 50% ethanol at RT for 1 min, and then replacing it with 1X PBS at RT for 1 min. The cells were permeabilized using 1X PBS + 0.1% Tween 20 (PBS/T) at RT for 10 MIN. The PBST was replaced by 1X PBS and washed once more in PBS. We applied 2–4 drops of hydrogen peroxide to each slide and incubated at RT for 10 min. Then removed the hydrogen peroxide solution and rinsed twice with PBS. We added the protease III solution (1:15 dilution in 1X PBS) for 10 min at RT. During this incubation, we warmed the probes for 10 min in a 40°C water bath and then allowed them to cool for 10 min. Following the protease III incubation, we washed the slides in distilled water twice, and then added the probes for 2 h in the RNAscope Hybridization oven at 40°C. After 2 hours, we gently washed the slides in 1x RNAscope wash buffer twice for 2 min at RT. After the probe hybridization, we added the Amp 1 and 2 solutions successively for 30 min in the RNAscope Hybridization oven at 40°C and then Amp 3 solution for 15 min. All of these steps were interspersed with 2 washes for 2 min in the 1x RNAscope wash buffer solution. We then added the HRP-C1 signal for 15 min in the RNAscope Hybridization oven at 40°C, followed by the diluted Opal Dye for signal development for 30 min in the RNAscope Hybridization oven at 40°C, and finally a 15 min incubation in the RNAscope Hybridization oven at 40°C with the HRP blocker. In between all of these steps, we again performed 2 washes for 2 min with the ACD wash buffer solution. We then repeated the HRP signal, the Opal Dye, and the HRP blocker for the C2 and C3 probes. We then mounted the slides by adding 4 drops of DAPI for 30 secs, removing it immediately and placed 1-2 drops of Prolong Gold antifade mounting media onto the slide. We carefully placed a 24 mm x 50 mm glass coverslip over the 18 mm round coverslips and dried the slides for atleast 30 min before imaging. Fluorescent images of cells were acquired with a Leica SP8 confocal scanning microscope equipped with a 100X oil-immersion objective.

### Preparation of OCT sections from brain microtissue

OCT embedding and sectioning were performed as previously described^58^. Briefly, samples were fixed in 4% PFA for approximately 30 minutes at room temperature (RT) and washed three times with DPBS. Samples were then incubated for 24 to 48 hours at 4°C in DPBS containing 30% sucrose, until the spheres had sunk. Next, samples were transferred to a 1:1 solution of OCT and DPBS containing 30% sucrose. They were then incubated in OCT for 30 minutes on ice before being embedded in OCT and frozen. OCT-embedded samples were cryosectioned at 20 µm intervals using a Leica cryostat, followed by the immunocytochemistry (ICC) protocol described below.

### Immunostaining for OCT sections from brain microtissue

Slides were washed once with DPBS and incubated with blocking buffer (5% BSA in PBS with 0.2% Triton X-100) for 1 hour at room temperature (RT). Sections were then incubated with primary antibodies diluted in 2% BSA in PBS with 0.1% Triton X-100 overnight at 4°C.

The following day, after washing with PBS, sections were incubated with secondary antibodies diluted in 2% BSA in PBS with 0.1% Triton X-100 for 1 hour at RT. Slides were then mounted using DAPI Fluoromount-G® (SouthernBiotech). Images were acquired using a ZEISS LSM980 confocal microscope.

### Whole-mount staining after clearing brain microtissue

Whole-mount staining after tissue clearing was performed based on previously published protocols^31,32^ with slight modifications. Samples were fixed with 4% paraformaldehyde (PFA) overnight at 4°C. After washing with DPBS, spheres were incubated with CUBIC-L (TCI, #T3740) overnight at 37°C. They were then washed with DPBS and incubated with primary antibodies diluted in 5% BSA in DPBS containing 0.3% Triton X-100 overnight at 37°C. After washing with DPBS, samples were incubated with secondary antibodies diluted in 5% BSA in DPBS with 0.3% Triton X-100 overnight at 37°C. The following day, samples were washed with DPBS and incubated with 1% formaldehyde in DPBS overnight at room temperature. Spheres were washed again with DPBS and transferred to a 96-well glass-bottom plate (Cellvis, #P96-1.5H-N) containing CUBIC-R (TCI, #T3741). Images were acquired using a ZEISS LSM980 confocal microscope.

### Computational pipeline for heuristic-based automated cell segmentation Overview of the Analysis Pipeline

Image processing and analysis for figures 6 (e, f), and S6 (d, e) were performed using a custom pipeline developed in Python 3 and available at https://github.com/chesnov/3D_microglia_segment.git. The graphical user interface was built using PyQt5 and Napari. The core computational pipeline relies on several open-source libraries, including NumPy for numerical operations, SciPy and Scikit-image for image processing algorithms, skan for skeleton analysis, and Pandas for data management.

The pipeline is designed to process both 3D volumetric images and 2D images through distinct but logically parallel workflows. For both 2D and 3D analysis, the process is structured into four sequential stages: (1) Initial Raw Segmentation, (2) Artifact Removal and Edge Trimming, (3) Soma-based ROI Refinement, and (4) Quantitative Feature Extraction. To handle large image datasets, the pipeline extensively uses memory-mapped files (numpy.memmap) for intermediate segmentation results, minimizing RAM usage. Computationally intensive tasks are accelerated using Python’s multiprocessing capabilities. All key parameters were made tunable through the graphic user interface and automatically exported to/loaded from a yaml file.

### 3D Segmentation and Analysis Pipeline

#### 1. Initial Raw Segmentation

The initial segmentation aims to generate a preliminary mask of all cellular structures in the 3D volume.

1. **Tubular Structure Enhancement:** To improve the detection of fine cellular processes, a slice-by-slice enhancement was applied. Each Z-plane of the input volume was independently processed using both Frangi and Sato vesselness filters (skimage.filters.frangi, skimage.filters.sato). The maximum response from these two filters was taken to create the final enhanced slice. This process was parallelized across Z-planes for efficiency. An optional initial 3D Gaussian smoothing could be applied before this step.
2. **Intensity Normalization:** The enhanced volume was normalized slice-by-slice along the Z-axis. For each plane, the intensity value at a high percentile (e.g., 98th) was mapped to a standardized value of 1.0, effectively balancing the brightness across different depths to compensate for light penetration artifacts at deeper imaging planes.
3. **Thresholding and Binarization:** A global intensity threshold was calculated from a random sampling of voxels in the normalized volume, corresponding to a user-defined lower percentile (e.g., 25th). All voxels with an intensity above this threshold were included in the initial binary mask.
4. **Fragment Connection:** To connect cellular processes that may have been broken during thresholding, an anisotropic morphological closing operation was performed (scipy.ndimage.binary_closing). The structuring element was dynamically shaped based on the physical voxel spacing to preferentially close gaps along axes with lower resolution.
5. **Small Object Removal:** The resulting binary mask was cleaned by removing all connected components smaller than a user-defined voxel count (skimage.morphology.remove_small_objects), eliminating noise and small, non-cellular fragments. The final output of this stage is an integer-labeled mask where each connected component has a unique ID.

### 2. Artifact Removal and Edge Trimming

This stage refines the raw segmentation by removing artifacts near the edges of the tissue.

1. **Tissue Hull Generation:** A 3D convex hull representing the overall tissue volume was generated. This was achieved by first creating a base tissue mask using Otsu’s thresholding method on the original intensity image. For each Z-slice, this tissue mask was combined with the raw cell segmentation mask, and a 2D convex hull was computed. These 2D hulls were stacked and smoothed in 3D using morphological closing and opening operations to create a coherent 3D tissue hull.
2. **Edge Trimming:** A boundary region of the hull was defined by eroding the smoothed 3D hull. Voxels within the raw segmentation that were located inside this boundary region *and* fell below a brightness cutoff (defined as a factor of the segmentation threshold from Step 1) were removed to compensate against the staining artifact at the edge of the tissue.
3. **Post-Trim Refinement:** Trimming can fragment objects at the tissue edge. A multi-step process was used to correct this:
⃘ **Relabeling:** Any object that was split into multiple disconnected components by the trimming was relabeled, with each new component receiving a new unique ID.
⃘ **Porosity Healing:** A novel healing step was applied to objects that were made porous by trimming. For each affected object, its 3D convex hull was calculated. Within this hull, any voxels that were part of the object *before* trimming but were removed, were restored. This effectively fills unnatural holes and concavities introduced by the trimming process.
⃘ **Merging:** Finally, any objects that became physically adjacent after the healing step were merged and assigned a single ID.

### 3. Soma Detection and Cell Separation

This stage identifies individual cells and separates any objects in the mask that contain multiple cell bodies (somas).

1. **Soma Candidate Detection:** A multi-heuristic approach was used to identify potential somas within each segmented object. The algorithm iteratively generated core candidates by thresholding each object based on both its Euclidean distance transform (using various ratios of the maximum distance) and its intensity distribution (using various percentiles).
2. **Candidate Filtering:** These candidates were rigorously filtered. True somas were identified from this pool based on a combination of criteria, including physical volume, thickness (maximum value in the distance transform), and aspect ratio, which was calculated via Principal Component Analysis (PCA) on the candidate’s voxel coordinates.
3. **Cell Separation:** Objects containing multiple valid somas were separated using a hybrid watershed and graph-based algorithm:

⃘ **Marker Generation:** For each multi-soma object, a graph was constructed where nodes were the detected somas. An edge was drawn between two somas if the intensity of the dimmest path between them in the original image (skimage.graph.route_through_array) was high relative to the soma intensities, suggesting they belong to the same cell. The connected components of this graph became the markers for the watershed algorithm.
⃘ **Guided Watershed:** A watershed transform (skimage.segmentation.watershed) was applied using these markers on a landscape derived from the negative distance transform of the object, optionally weighted by image intensity.
⃘ **Interface-Based Merging:** The initial watershed separation was refined by analyzing the interface between the resulting segments. A graph was built where nodes were the new segments. Edges were weighted based on metrics at the interface, including the mean intensity relative to the somas and the local intensity difference. If the segment brightness was too similar on both sides of the separation, or if their connection intensity was too bright, the segments were merged back together. The final output of this stage is a segmentation mask where each label corresponds to a single cell.

### 4. Quantitative Feature Extraction

A comprehensive set of features was calculated for each finally segmented 3D cell.

- **Morphological and Positional Features:** Basic properties including volume (µm³), surface area (µm²), bounding box volume, and sphericity were calculated for each cell. The median depth of each cell along the Z-axis was also computed. These calculations were optimized by processing only the bounding box region of each cell.
- **Proximity Analysis:** The shortest Euclidean distance between the surfaces of every pair of cells was calculated. This was achieved by extracting the surface voxels for each object, building a k-d tree (scipy.spatial.cKDTree) for one object’s surface points, and querying it to find the nearest neighbor for every point on the other object’s surface. The coordinates of the two closest points were also recorded.
- **Ramification Analysis:** The complexity of each cell was quantified via skeletonization.

⃘ A 3D morphological skeleton was generated for each cell (skimage.morphology.skeletonize).
⃘ An optional length-based spur pruning was performed, removing short, terminal branches smaller than a specified physical length.
⃘ The skan library was used to analyze the skeleton’s graph structure and calculate the total skeleton length.
⃘ To obtain a more biologically accurate measure of ramification, a custom topological analysis was performed on the skeleton. The number of true endpoints (pixels with 1 neighbor), true junctions (pixels with ≥3 neighbors), and true branches were calculated from the pixel-wise connectivity, correcting for computational artifacts inherent in standard skeleton analysis.

### 2D Segmentation and Analysis Pipeline

The 2D pipeline mirrors the four-stage logic of its 3D counterpart, with algorithms adapted for 2D images.

- **Initial Segmentation & Artifact Removal:** The 2D pipeline performs the same sequence of enhancement (Frangi/Sato on the 2D image), normalization, thresholding, fragment connection, and small object removal. Edge trimming is also performed by generating and using a 2D convex hull of the tissue. All morphological operations use 2D structuring elements (e.g., skimage.morphology.disk).
- **Soma Detection and Cell Separation:** The process for identifying and separating multi-soma cells is identical in principle to the 3D version, but all operations—including distance transforms, PCA for aspect ratio, watershed, and interface analysis—are performed in 2D.
- **Proximity:** Shortest distance between the boundaries of 2D cells was calculated using k-d trees on the boundary pixels.

### Time-lapse imaging for 3D brain microtissue (Figure 6G)

Day 16 tri-lineage brain microtissue containing membrane-tdTomato-labeled microglia was replated onto a Matrigel-coated 35 mm dish with a No. 1.5 coverslip (MatTek). The following day, time-lapse imaging was performed using ZEISS Lattice Lightsheet 7 (Zeiss) with 44× objective over approximately 4 hours at 1-minute intervals. Images were cropped and deconvoluted using ZEN software, and a movie was created using ImageJ.

### α-Synuclein Preformed Fibril (PFF) Preparation

α-Syn PFFs were prepared by Meredith Jackrel as previously described^62^. Briefly, monomeric α-syn was filtered through a 0.2µM filter and diluted in fibrillization buffer (pH 8 20mM Tris, 100 mM NaCl), then incubated at 37°C with agitation at 1500rpm for 7 days to produce fibrils. The resulting preparation was then centrifuged at 21130×g for 30 minutes at room temperature, after which the supernatant was removed, and a BCA assay was performed to quantify the concentration of the resulting fibrils. PFFs were then resuspended to 5 mg/mL in fibrillization buffer, snap frozen, and stored at-80°C until use. For use in experiments, frozen aliquots of PFFs were thawed at room temperature, diluted to 25µM in fibrillization buffer, then sonicated in a cup horn sonicator (Sonics) at an amplitude of 65 for 3 minutes (30s on/30s off). Once thawed and sonicated, the fibrils were immediately added to cell cultures.

### Data availability

Raw RNA-seq sequencing and normalized expression file has been deposited at SRA/GEO and are publicly available as of the date of publication (accession number: SRA: PRJNA1285605, GEO: GSE301787).

